# Room-temperature light-state structure of the AsLOV2-Rac1 optogenetic actuator by sheet-on-sheet serial femtosecond crystallography

**DOI:** 10.64898/2026.10.27.762267

**Authors:** Magdalena Pachota, Magdalena Wiśniewska, Madan Kumar Shankar, Martin V. Appleby, Anna Koczurowska, Ayesha Asghar, Michał Kępa, Ran Ke, Aleksandra Solecka, Teresa Durán Kobylecka, Bence Olasz, Sandra Mous, John H. Beale, Guillaume Gotthard, Przemyslaw Nogly

## Abstract

**Synopsis:** Serial femtosecond crystallography on sheet-on-sheet chips gives, to our knowledge, the first light-state structure of the optogenetic actuator AsLOV2-Rac1, recorded at room temperature with about 80 % FMN-cysteine adduct free of X-ray radiolysis. Apart from the poorly ordered N-terminal residue, light-induced Cα displacements above 0.3 Å are confined to the chromophore environment, the Dα helix and the Gβ-Hβ and Hβ-Iβ loops of LOV2; in the crystal, the Jα helix stays folded and no local change is detected in the Rac1 domain.

Light-oxygen-voltage (LOV) domains are blue-light sensors that make fused effector proteins optically controllable. The AsLOV2-Rac1 fusion (PA-Rac1) is one of the most widely used of these tools, yet the structural basis of its activation has been inferred largely from cryogenic structures and from solution studies, mostly of the isolated AsLOV2 domain. Here, room-temperature dark and light-state structures of AsLOV2-Rac1 at 1.8 Å resolution are reported, determined by serial femtosecond crystallography at the SwissFEL Cristallina beamline with sheet-on-sheet (SOS) chip sample delivery. Repeated 450 nm pump-laser excitation during chip rastering accumulated the FMN-cysteine adduct in about 80 % of the molecules, giving strong Fourier difference maps. To our knowledge, this is the first light-state structure of PA-Rac1 or of any AsLOV2 fusion and the first AsLOV2 light state in which the thioether adduct is not compromised by X-ray radiolysis. Apart from the N-terminal residue and one marginal Rac1 loop residue, Cα displacements above 0.3 Å between dark and light models recorded at the same temperature are confined to the chromophore environment, the Dα helix and the Gβ-Hβ and Hβ-Iβ loops of AsLOV2. These changes, including the Ile445 rotamer switch, are also seen in light-state structures of the isolated AsLOV2 domain, but the Asn414 rotation reported there is absent in the fusion. Comparison with the cryogenic dark-state structure also shows temperature-dependent differences that exaggerate the apparent changes. In the crystal, the GTP, the Mg^2+^ site and the switch regions of Rac1 show no detectable light-induced change and the Jα helix does not unfold, although transient or low-population states cannot be excluded. This modest response is consistent with the small difference in effector affinity between dark- and lit-state mutants in solution, with lattice contacts and an interdomain Ca^2+^ ion as possible further contributions.

## Introduction

Phototropins are plant blue-light photoreceptors that regulate phototropism, the rapid inhibition of hypocotyl growth, chloroplast migration and stomatal opening (Lin, 2002; Sakai *et al*., 2001; Kinoshita *et al*., 2001; Folta & Spalding, 2001). Each phototropin carries two light-oxygen-voltage (LOV) domains, LOV1 and LOV2, and a C-terminal serine/threonine kinase. Blue light triggers a photocycle in which a covalent adduct forms between the C4a atom of the flavin mononucleotide (FMN) chromophore and a conserved cysteine, accompanied by a blue shift of the absorption (Salomon *et al*., 2000; Halavaty & Moffat, 2007). The resulting structural changes spread from the FMN pocket through the domain core to the C-terminal Jα helix and on to the kinase (Halavaty & Moffat, 2007).

LOV domains control a wide range of effector domains in nature (Glantz *et al*., 2016), which makes them attractive modules for engineering light-controlled proteins (Flores-Ibarra *et al*., 2024). The LOV2 domain of *Avena sativa* phototropin 1 (AsLOV2) has been fused to dihydrofolate reductase (Lee *et al*., 2008) and to the diaphanous autoregulatory domain of the formin mDia1, which enabled the optogenetic activation of endogenous diaphanous-related formins (Rao *et al*., 2013); a similar fusion was used to trigger actin polymerization in the nucleus (Baarlink *et al*., 2013; Plessner & Grosse, 2019). It is also the light-sensing part of tunable light-inducible dimerization tags (TULIPs), used to recruit proteins to cellular structures and to control the transport of endosomes and mitochondria (Strickland *et al*., 2012; van Bergeijk *et al*., 2015), of a light-inducible nuclear localization signal (LINuS) (Niopek *et al*., 2014), and of LOVTRAP, which reversibly sequesters proteins at mitochondria (Wang *et al*., 2016).

In photoactivatable Rac1 (PA-Rac1), AsLOV2 is fused to the N terminus of Rac1, a Rho-family GTPase that controls actin cytoskeletal dynamics, cell growth, endocytosis and the generation of reactive oxygen species, and whose aberrant activity has been linked to cancer and to neurodegenerative and cardiovascular diseases (Bosco *et al*., 2009). In the dark, the LOV2 domain sterically blocks effector binding to Rac1. Light is thought to unwind the Jα helix between the domains and to release Rac1, which in its GTP-bound state binds effectors such as p21-activated kinase 1 (PAK1) (Bokoch, 2003; Bishop & Hall, 2000). Because the activating light can be confined to part of a cell, PA-Rac1 controls Rac1 activity in space as well as in time, and hence cell motility and morphology in living cells (Wu *et al*., 2009). It remains widely used, for example in studies of lamellipodia formation by cryo-electron tomography (Inaba *et al*., 2025), of Rac1 signalling in the long-term synaptic plasticity of dendritic spines (Saneyoshi *et al*., 2025) and of the Rho GTPase-effector ensemble that governs spontaneous cell migration (Lee *et al*., 2025).

Cryogenic dark-state structures of PA-Rac1 and of its C450A and C450M variants (PDB entries 2WKP, 2WKQ and 2WKR, at 1.9, 1.6 and 2.2 Å resolution) show that the LOV2 domain occludes the effector-binding surface of Rac1 and that Jα is folded as in the isolated AsLOV2 domain (Wu *et al*., 2009). In solution, hydrogen-deuterium exchange and small-angle X-ray scattering indicate that light does not fully release Rac1 from the LOV2 cage but shifts a population of pre-existing conformations, with caged and unstructured Jα states co-existing even in the dark (Winkler *et al*., 2015). To our knowledge, no light-state structure of PA-Rac1 or of any other AsLOV2 fusion has been reported, and X-ray radiolysis breaks part of the thioether adduct in the synchrotron light states of the isolated AsLOV2 domain (Halavaty & Moffat, 2007; Dietler *et al*., 2022).

Here we report room-temperature dark and light-state structures of PA-Rac1 determined by serial femtosecond crystallography (SFX) at the Cristallina beamline of SwissFEL, using sheet-on-sheet (SOS) chips (Doak *et al*., 2024) to deliver fragments of large rod-shaped crystals. Repeated pump-laser excitation drove about 80 % of the molecules into the adduct state. Because SFX data are recorded before radiation damage develops, the adduct is not compromised by X-ray radiolysis. By comparing dark and light structures recorded at the same temperature, we ask which parts of the light response of isolated AsLOV2 operate within the fusion, and whether the Jα helix and Rac1 respond in the crystal.

## Materials and methods

### Protein expression and purification

The expression and purification protocol was adapted from Wu et al. (2009). A codon-optimized construct encompassing residues 404-546 of *Avena sativa* phototropin 1 and 4-180 of human Rac1, carrying the Q61L, E91H and N92H substitutions of the original PA-Rac1 design (Wu *et al*., 2009), was ordered in the pQE-60 plasmid from GenScript, with a stop codon before the vector-encoded C-terminal His tag, and transformed into *E. coli* XL10-Gold. The construct carries an N-terminal MGGSHHHHHHHHGS tag, which was not removed; the tag is disordered in both structures, and the full sequence is given in the Supporting information. Cultures were grown in LB medium to OD_600=_ 0.6-0.8, and expression was induced with 1 mM isopropyl β-D-1-thiogalactopyranoside (IPTG) at 27 °C overnight. Cells were harvested by centrifugation (20 min, 6000 × g, 4 °C) and stored at −80 °C. Thawed cells were resuspended in 20 mM Tris pH 8.5, 50 mM NaCl, 5 mM MgCl_2,_ 2 mM β-mercaptoethanol and 40 µM GTP and disrupted in a continuous-flow cell homogenizer at 159 MPa (23 kpsi). The lysate was cleared by centrifugation (1 h, 30 000 × g, 4 °C) and applied to a HisTrap HP column; the His-tagged protein was eluted with an imidazole gradient and dialyzed overnight at 4 °C against 10 mM Tris pH 8.5, 20 mM NaCl, 5 mM MgCl_2,_ 2 mM β-mercaptoethanol and 40 µM GTP. The protein was then purified on a Capto HiRes Q 5/50 column with a NaCl gradient and on a HiLoad Superdex 75 pg column in 10 mM Tris pH 8.5, 20 mM NaCl, 5 mM MgCl_2,_ 2 mM DTT and 40 µM GTP. GTP was included in all buffers to obtain a more homogeneous protein. The protein was concentrated to 18 mg ml⁻¹, aliquoted and snap-frozen in liquid nitrogen.

### Crystallization

The crystallization conditions deposited by Wu et al. (2009) with PDB entry 2WKP were adapted to batch crystallization to produce the large amounts of crystals needed for serial crystallography. The protein (18 mg ml⁻¹) was added drop by drop, at a 2:1 protein:precipitant (v/v) ratio, to a precipitant solution containing 18 % (w/v) polyethylene glycol (PEG) 8000, 100 mM sodium cacodylate pH 5.5 and 150 mM calcium acetate, with gentle mixing. The tubes were incubated in the dark at 20 °C, and rod-shaped crystals 20-300 µm long appeared after 24 h and sedimented.

### Data collection and processing

Diffraction data were collected at the Cristallina beamline of SwissFEL using SOS chips with 6 µm Mylar films (Doak *et al*., 2024). For chip loading, most of the precipitant above the sedimented crystals was removed, and the dense slurry was mixed 1:1 with 5 % hydroxyethyl cellulose in precipitant solution, which also broke the crystals into smaller fragments. The X-ray beam size was 2.2 × 3.6 µm (FWHM) and the transmission was set to 40 %. Dark and illuminated datasets were collected from separate SOS chips, rastered in 25 µm steps horizontally and vertically. For the illuminated dataset, the crystals were excited with a 450 nm, 3 ns (FWHM) pump laser (EKSPLA; 53 × 53 µm spot at 1/e², 3.63 mJ/cm²) synchronized to the SwissFEL timing system, with the pump pulse preceding the X-ray pulse by 100 ns. Because the SOS chip is a sandwich of two transparent films without wells between raster positions, the pump spot is about twice the raster step and much larger than the X-ray focus, and the sample scatters the pump light, each probed crystal volume had received a series of pump pulses as the chip moved through the beam, over the seconds preceding its exposure. This is short compared with the thermal recovery of the photoadduct, which has a time constant of about 80 s in AsLOV2 at 22 °C (Zayner *et al*., 2012) and a half-life of 43 s in PA-Rac1 at room temperature (Wu *et al*., 2009). The illuminated dataset therefore samples a light state accumulated over repeated pump pulses rather than the transient state 100 ns after a single pulse; the experiment is not time-resolved. A pump spot of this size reaches crystals at neighbouring positions in transparent fixed targets (Gotthard, Flores-Ibarra *et al*., 2024), and pump-probe measurements on SOS chips that must avoid this focus the pump to about 6 µm (Barends *et al*., 2026). We use the term light state for the structure of the illuminated crystals, in which most molecules carry the adduct; the term does not imply that the signalling state, with an unfolded Jα helix, is reached.

The diffraction images were processed with CrystFEL (White *et al*., 2012), using the peakfinder8 (Barty *et al*., 2014) and xgandalf (Gevorkov *et al*., 2019) algorithms for peak finding and indexing, respectively, with --threshold=10, --min-snr=4 and --min-pix-count=2. The indexing ambiguity was resolved with ambigator (Brehm & Diederichs, 2014; White *et al*., 2016) with the options -y 321_H and --operator=-h,-k,l. The reflection intensities were scaled and merged with partialator (--model=unity), and data statistics were calculated with check_hkl and compare_hkl.

### Structure determination and refinement

Molecular replacement was performed in Phaser (McCoy *et al*., 2007) with the cryogenic PA-Rac1 structure (PDB entry 2WKP) as search model (Wu *et al*., 2009). The dark structure was built and refined at 1.8 Å with Coot (Emsley *et al*., 2010) and PHENIX (Liebschner *et al*., 2019), using a common set of free-R reflections for the dark and light datasets. Selected comparative refinements with REFMAC5 (Murshudov *et al*., 2011) were used to test robustness to the refinement protocol. The light-state structure was refined against the light data starting from the dark model. The Cys450-FMN region was described by three alternate conformations, each comprising a Cys450 conformer and its FMN: the covalent adduct between Cys450 Sγ and FMN C4a, and the two dark-state Cys450 conformers with non-covalently bound FMN. Their occupancies, refined as one group constrained to sum to one, were 0.78 for the adduct and 0.10 and 0.12 for the two dark-state conformers, whereas the same refinement against the dark data returned no adduct (≤0.02). Residues with ambiguous difference density were inspected with alternative rotamers and modelled conservatively where no single light-specific conformation was supported. Because the amide oxygen and nitrogen of Gln513 cannot be told apart directly at this resolution, both orientations were refined and compared with density tests calibrated on asparagine and glutamine amides of known orientation. These tests and a two-conformer test of Asn414 are described in the Supporting information (Figures S1-S3).

Model quality was assessed with MolProbity (Williams *et al*., 2018): 98.8 % (dark) and 97.0 % (light) of the residues lie in the favoured regions of the Ramachandran plot, with no Ramachandran outliers. Electron density was interpretable for residues 404-723 in both structures. Cys648 (Rac1 Cys105) carries a covalent modification in both datasets, modelled as S-sulfocysteine with refined group occupancies of about 0.7 (dark) and 0.66 (light); its identity remains to be confirmed by mass spectrometry (Supporting information).

### Difference maps

Light-minus-dark F_o-_F_o m_aps were calculated with phenix.fobs_minus_fobs_map (Liebschner *et al*., 2019) using the multiscale option, a resolution range of 1.8-30 Å, a sigma cutoff of 3.0 and phases from the dark model, and were inspected in Coot (Emsley *et al*., 2010). Unless stated otherwise, difference-map values quoted below are local maxima or minima next to the named residue, and σ is the root-mean-square density of the map over the unit cell.

### Displacement and rigid-body analyses

For the displacement analysis, the models were superposed in PyMOL (Schrödinger, LLC, 2015) separately for each domain on Cα atoms (LOV2 core, residues 404-521; Rac1, residues 547-718), and Cα displacements, using the closest pair of conformers for residues with alternate conformations, were compared with a threshold of 0.30 Å. Published structures of the isolated AsLOV2 domain and of PA-Rac1 (Halavaty & Moffat, 2007; Dietler *et al*., 2022; Wu *et al*., 2009) were re-analysed with a common protocol, and rigid-body rotations of LOV2 relative to Rac1 were fitted to isomorphous difference data calculated with servalcat (Yamashita *et al*., 2021, 2023) and gemmi (Wojdyr, 2022); both analyses are described in the Supporting information.

## Results and discussion

### Light-state data from fragmented crystals on SOS chips

PA-Rac1 did not form homogeneous microcrystals but grew as large, irregular rods. SOS chips, which have no patterned wells and accept crystals of a broad range of sizes and size distributions (Doak *et al*., 2018; Gorel *et al*., 2025), allowed these rods to be crushed and loaded directly as a microcrystalline slurry with only a thin layer of sample per chip. Although diffraction extended to 1.58 Å, both datasets were truncated at 1.80 Å on the basis of CC-based criteria, completeness and the stability of R_free (_Table 1). Repeated excitation during rastering accumulated the adduct in about 80 % of the molecules. The light-state data therefore describe this accumulated state rather than a time-resolved snapshot, and they give strong and detailed difference maps. The combination of SOS-chip delivery, repeated pump excitation and matched-temperature comparison should be applicable to other slow-cycling LOV fusions, such as LINuS, TULIPs and LOVTRAP, and to other photoreceptors that form heterogeneous crystals.

**Table 1.** Data collection and refinement statistics.

|  | Dark | Light |
| --- | --- | --- |
| Beamline | SwissFEL Cristallina |  |
| Photon energy (keV) | 12.03 |  |
| X-ray pulse length (fs) | 25 (FWHM) |  |
| Repetition rate (Hz) | 100 |  |
| Detector | JUNGFRAU 8M |  |
| Temperature (K) | 298 |  |
| Sample delivery | SOS chips (6 $\mu\text{m}$ Mylar), 25 × 25 $\mu\text{m}$ raster | |
| Pump laser | None | 450 nm, 3 ns (FWHM), 53 × 53 $\mu\text{m}$ (1/e<sup>2</sup>), 3.63 mJ/cm<sup>2</sup>, 100 ns before the X-ray pulse |
| X-ray beam size ($\mu\text{m}$, FWHM) | 2.2 × 3.6 | 2.2 × 3.6 |
| Crystal size ($\mu\text{m}$) | 20-300 | 20-300 |
| Space group | $P3_221$ | $P3_221$ |
| Lattice parameters ($\text{\AA}$, $^\circ$) | 115.56, 115.56, 69.93; 90, 90, 120 | 115.56, 115.56, 69.93; 90, 90, 120 |
| Indexed patterns | 77,475 | 75,136 |
| No. of collected images | 294,440 | 294,616 |
| Indexing rate (% of collected images) | 26.3 | 25.5 |
| Resolution range ($\text{\AA}$) | 44.51-1.80 (1.86-1.80) | 40.66-1.80 (1.86-1.80) |
| No. of measurements | 66,495,539 | 55,308,635 |
| Unique reflections | 96,608 | 96,608 |
| Unique reflections, Friedel pairs merged | 50,044 | 50,026 |
| Multiplicity | 688.3 (463.3) | 572.5 (384.3) |
| Completeness (%) | 100.0 (100.0) | 100.0 (100.0) |
| Mean $I/\sigma(I)$ | 8.26 (1.91) | 6.71 (1.05) |
| CC* | 0.9941 (0.9317) | 0.9928 (0.8268) |
| CC$\frac{1}{2}$ | 0.9769 (0.7670) | 0.9718 (0.5193) |
| R-split (%) | 11.29 (51.24) | 12.76 (90.96) |
| Wilson B factor ($\text{\AA}^2$) | 31.3 | 32.0 |
| Refinement resolution ($\text{\AA}$) | 30.11-1.80 (1.83-1.80) | 30.11-1.80 (1.83-1.80) |
| Reflections (refinement) | 50,026 (2,718) | 50,009 (2,717) |
| Reflections, test set | 2,452 (4.9 %) | 2,451 (4.9 %) |
| R-work / R-free (%) | 15.49 / 17.54 (32.40 / 38.73) | 15.80 / 18.31 (31.14 / 35.48) |
| No. of atoms | 3013 (protein 2692, water 256, other 65) | 2975 (protein 2631, water 217, other 127) |
| B-factors ($\text{\AA}^2$, avg) | protein 35.39, water 49.54, other 27.30 | protein 41.78, water 53.57, other 30.62 |
| Ramachandran favoured (%) | 98.83 | 96.99 |
| Ramachandran allowed (%) | 1.17 | 3.01 |
| Ramachandran outliers (%) | 0.00 | 0.00 |
| RM SD bond lengths ($\text{\AA}$) | 0.007 | 0.011 |
| RM SD bond angles ($^\circ$) | 0.915 | 1.205 |
| PDB code | 35HK | 35HL |

\*Values in parentheses are for the highest-resolution shell. Unique reflections are counted with Friedel pairs unmerged (point group 321).

### Overall structure

The two domains are arranged as in the cryogenic structure, with the LOV2 module shielding the effector-binding surface of Rac1 (Figures 1 and 2). Structural alignment with PDBeFold (Krissinel & Henrick, 2004) gives a Cα RMSD of 0.25 Å to the cryogenic dark structure of Wu et al. (2009) (Figure 1(a)), with the largest deviations in the Hβ-Iβ loop, the Pro572-Gly573 loop next to the GTP site and the C terminus, which likely reflect the weaker density in these regions. Our dark and light structures differ by a Cα RMSD of only 0.18 Å. AsLOV2 adopts the PAS fold, flanked by an N-terminal cap (Leu404-Asn414) containing the AZα helix and by the Iβ-Jα loop (Thr517-Arg521) and the Jα helix (Asp522-Lys544), which interacts with the effector domain. Compared with the isolated AsLOV2 domain, the N-terminal cap is rotated at Leu408 and oriented towards Jα, a change induced by the adjacent Rac1 domain (Figure 1(b)). Rac1 contributes interface residues mainly from β1-β3, α1 and η2, and AsLOV2 from the Aβ-Bβ, Hβ-Iβ and Dα-Eα loops, the N-terminal cap and the C-terminal part of Jα (Figure 2 and Figure S4).

**Figure 1.**
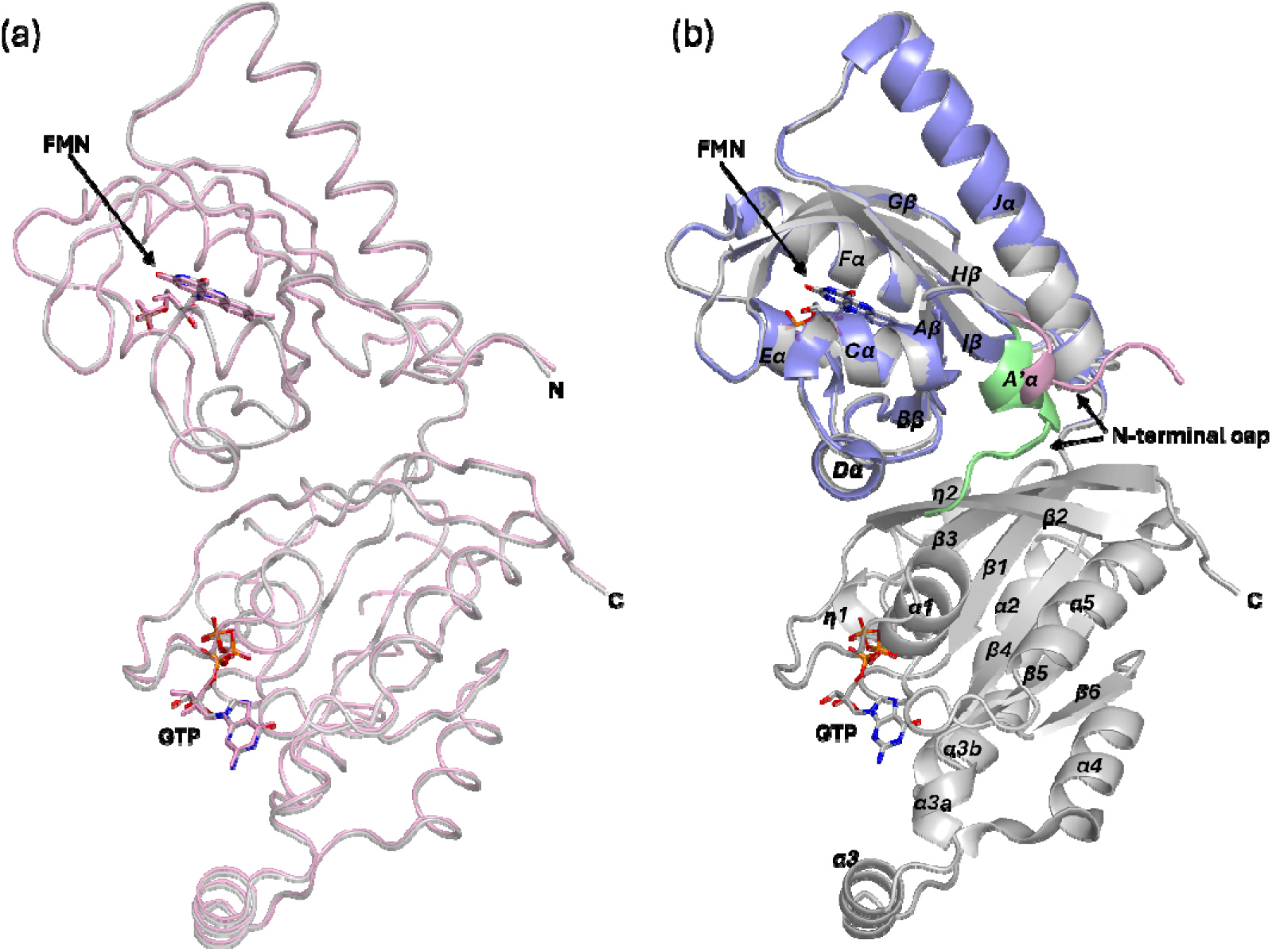
Cartoon representation of the AsLOV2-Rac1 structures. (a) Superposition of dark (grey) and the published AsLOV2-Rac1 structure (PDB: 2WKP, pink, cryo). (b) Superposition of dark-state structures of AsLOV2-Rac1 (grey) and the isolated AsLOV2 domain (PDB: 2V1A, slate, RT). N-terminal cap is marked in pink (AsLOV2-Rac1) and green (AsLOV2).

**Figure 2.**
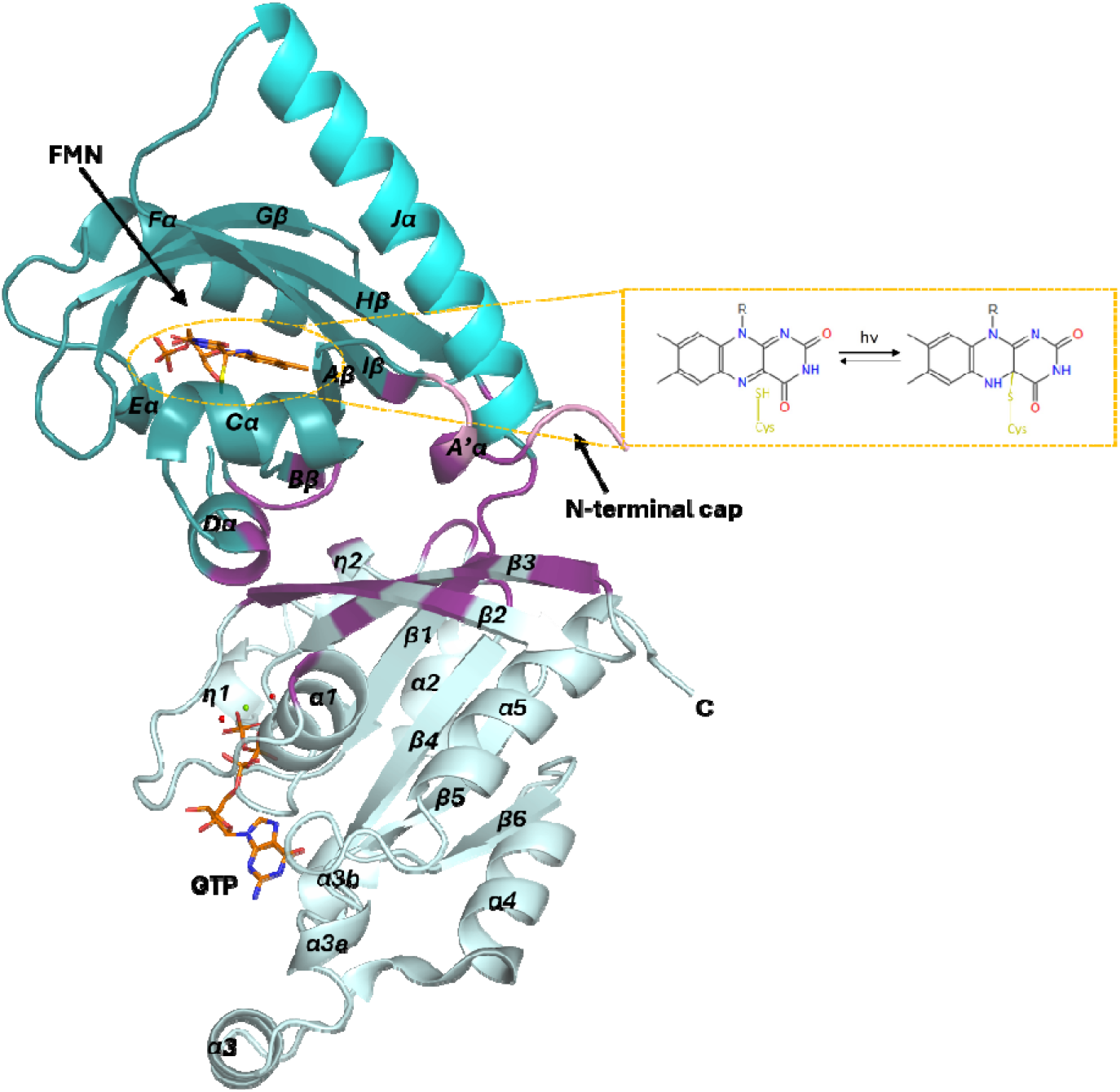
Cartoon representation of the AsLOV2-Rac1 structure in the light state. The AsLOV2 domain (deep teal) consists of a five-stranded antiparallel β-sheet where the FMN chromophore binding pocket is located (orange sticks), an N-terminal cap with an A□α-helix (pink), and the C-terminal Jα-helix that interacts with effector proteins (cyan). The sulfur of the conserved LOV Cys residue is pictured attached covalently to C4a of FMN with a yellow adduct bond. Rac1 (pale cyan) consists of one central β-sheet made up of six strands, five parallel and one anti-parallel, six α-helices and two short 3_10 h_elices. The nucleotide-binding pocket is occupied by GTP (orange sticks) and a magnesium ion (green sphere). Coordinated water molecules are shown as red spheres. Interface residues are marked in purple. The inset shows the light-induced formation of the covalent bond between the Cys450 thiol and FMN C4a.

GTP is bound in its canonical ground state. The γ-phosphate is fully occupied, and the Mg^2+^ ion is octahedrally coordinated by the hydroxyls of Thr560 and Thr578 (Rac1 Thr17 and Thr35), one oxygen each of the β-and γ-phosphates and two water molecules, with a water in line with the Pγ-O3β bond at 3.5 Å from Pγ. This is consistent with the GTPase-deficient Q61L background of PA-Rac1 (Wu *et al*., 2009), and neither dataset shows GTP hydrolysis. A Ca^2+^ ion from the crystallization medium bridges the two domains within the molecule. It is coordinated by the main-chain carbonyls of Glu443 and Leu446 in the Dα-Eα segment of AsLOV2, both carboxylate oxygens of Asp581 (Rac1 Asp38, switch I) and three water molecules. The same site is occupied in the cryogenic structure (PDB 2WKP), and calcium binding, with a dissociation constant of about 0.46 mM, stabilizes the sensor-effector interface and modulates the light response of PA-Rac1 in solution (Winkler *et al*., 2015).

### The photoadduct and the chromophore pocket

Light-minus-dark difference maps show strong features mostly in the LOV2 domain (Figure 3). The strongest are a +19.2 σ peak between Cys450 and FMN C4a and a −13.8 σ peak at a distal Cys450 conformation, consistent with formation of the cysteinyl-flavin thioether adduct. As in the isolated domain, Cys450 adopts two conformations in the dark and forms the adduct in the light state (Figure 4(a) and 4(b)), rotating by about 105° around the Cα-Cβ bond. Paired peaks of about 10 σ above and below the isoalloxazine ring reflect its tetrahedral distortion and an asymmetric displacement towards the cysteine side, with the ring plane tilted by about 7°, whereas the ribityl chain and phosphate barely move (Figure 4(c)). Apart from Gln513 (below), the hydrogen bonds between the isoalloxazine ring and the surrounding amide side chains are largely conserved; Gln454, Asn482 and Asn492 show difference features consistent with shifts of 0.2-0.3 Å, close to the coordinate error, which accommodate the displaced flavin ring (Figure S5). Polder omit maps (Liebschner *et al*., 2017) show continuous density for the adduct and only weak residual features at the dark-state position.

**Figure 3.**
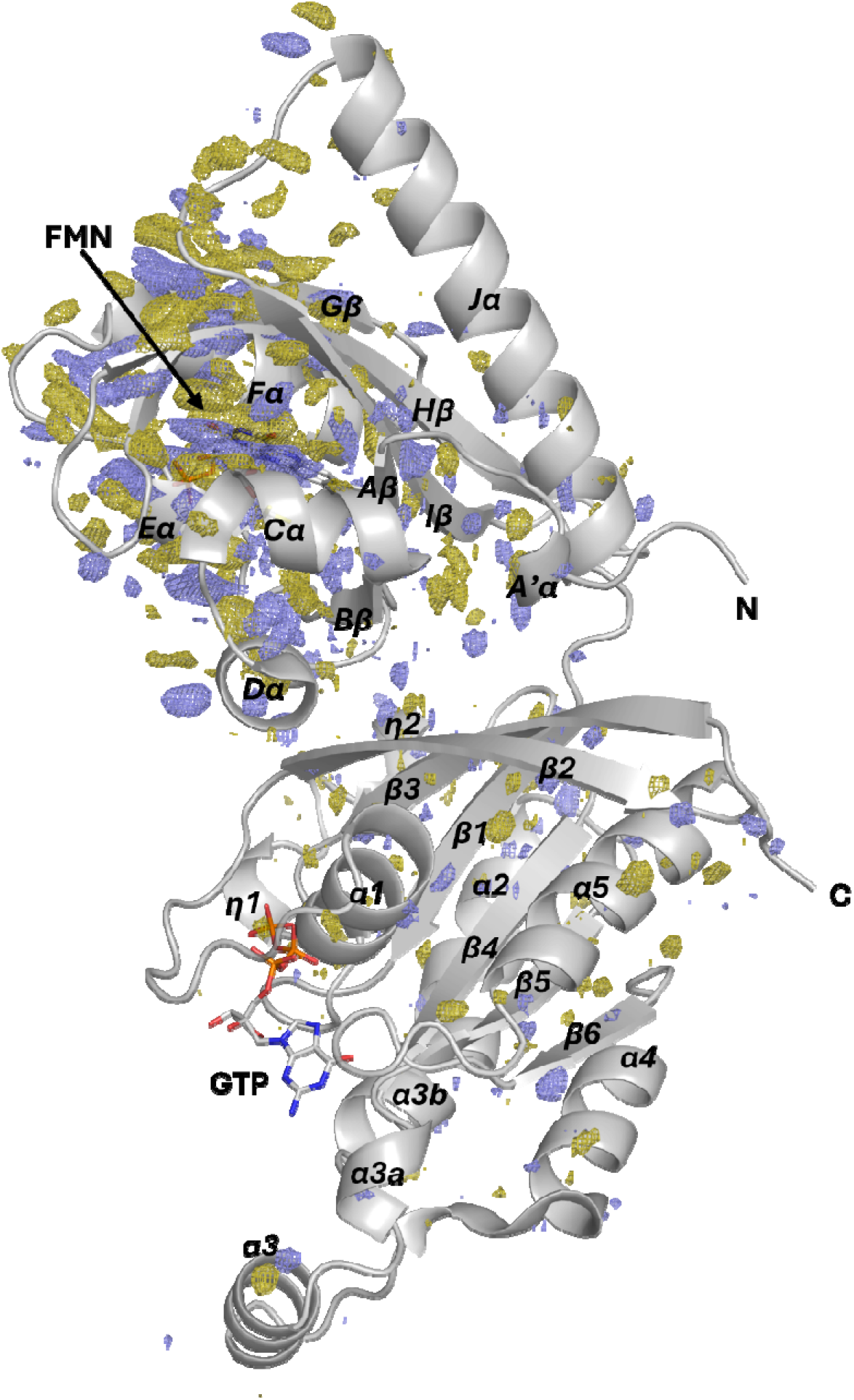
Overall light-induced structural changes in AsLOV2-Rac1. The F_light_ − F_dark_ difference electron density map is contoured at ±3.0 σ, with positive and negative features displayed as blue and yellow meshes, respectively.

**Figure 4.**
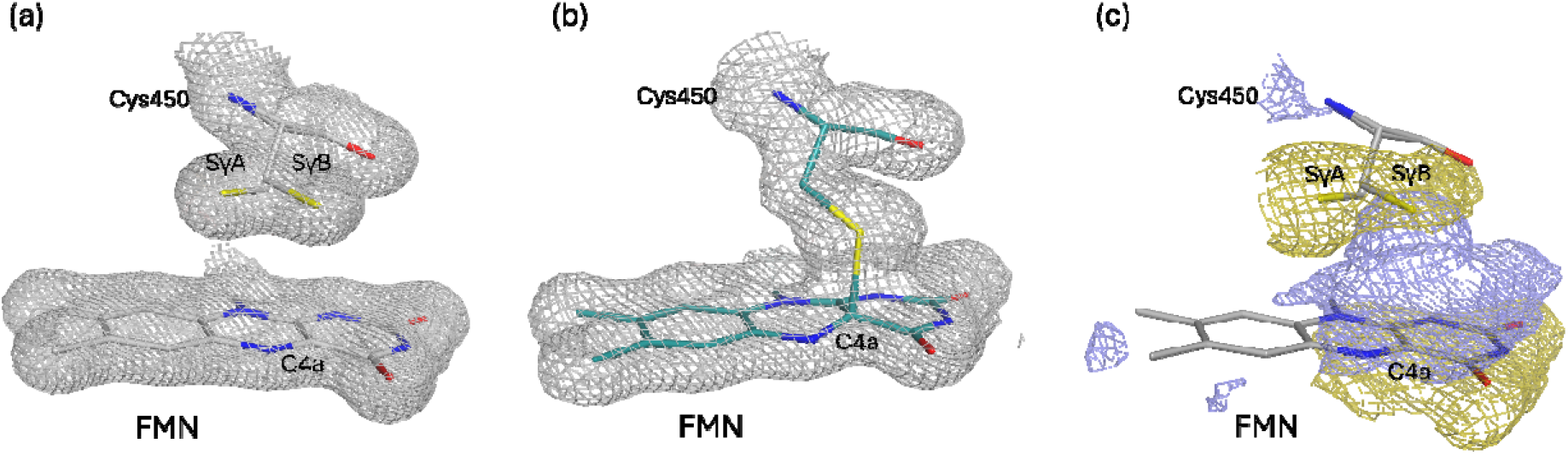
Electron density maps of the FMN and Cys450 in the dark and light states of AsLOV2-Rac1. The 2F_o_ − F_c_ map of the dark state structure (a) and the light state (b) contoured at 1.0 σ. The photoactive Cys450 exhibits two conformations in the dark state. Upon blue-light exposure, the side chain of Cys450 rotates and the thiol forms a covalent bond with C4a of the FMN chromophore. In comparison with the dark state, the plane of the isoalloxazine ring is tilted by about 7°, causing a slight shift of the amino acids surrounding the binding pocket (c). The difference electron density map F_light_ − F_dark_ contoured at ±3.0 σ, with positive and negative features shown as blue and yellow meshes, respectively.

About 20 % of the molecules in the light data carry no adduct, and in them Cys450 occupies both dark-state rotamers. We attribute this fraction to residual dark-state molecules, consistent with the incomplete photoactivation proposed for the 20 % dark-state population retained in the light state of isolated AsLOV2 (Dietler *et al*., 2022). It does not resemble the pseudo-ground state DZ described for illuminated crystals of the LOV2 domain of Arabidopsis thaliana phototropin 2, which has a single cysteine conformation and a disordered C terminus (Aumonier *et al*., 2022). Our residual population keeps both cysteine rotamers, and Jα remains ordered.

Because SFX data are recorded before radiation damage develops, the adduct is not affected by the X-ray radiolysis of the thioether bond seen in synchrotron light states of LOV2 (Halavaty & Moffat, 2007; Dietler *et al*., 2022). In σ_A-_weighted maps recalculated from the deposited data, the density at the midpoint of the Cys450 S-C4a bond, relative to the mean density of the flavin ring, is 1.01 in our light state, similar to the midpoints of the covalent bonds within the flavin ring (0.9-1.2 in our maps). It is 0.71 and 0.44 in the synchrotron light states 2V1B and 2V0W of the isolated domain and 0.03 in 7PGY, where no adduct was modelled, and it is negative at the same position in our dark map, where there is no bond.

### The conserved LOV2 light response operates in the fusion

Beyond the chromophore pocket, the strongest features lie in the main chain of strand Gβ, at Val478 (+5.0/-5.6 σ), Gln479 (+5.5/-7.0 σ), Leu480 (+6.9/-5.2 σ) and Ile481 (+6.9/-8.8 σ), and in the side chains of Phe452 (+5.6/-7.0 σ), Leu453 (+5.4/-5.8 σ) and Trp491 (+5.9/-6.0 σ) (Figure 5). Because peak heights are relative measures, we also give the atomic shifts after superposition on the LOV core: at most 0.1-0.2 Å in the Gβ main chain and in Trp491, below the coordinate error, and 0.6 and 0.7 Å in the side chains of Phe452 and Leu453. On helix Dα, Ile445 (+6.0/-7.0 σ) adopts the rotamer of the cryo-trapped light state of isolated AsLOV2 (PDB 2V0W), switching χ1 from about +60° to about −68° and moving its side-chain atoms by up to 3.4 Å, and the preceding Glu444 shows paired features of +4.7/-4.4 σ (main-chain shift up to 0.7 Å). After superposition on the LOV core, Glu444 and the side chains of Ile427, Leu435, Cys450, Phe452 and Leu453 shift in the same directions as in the isolated-domain light states 2V0W and 7PGY (vector cosines 0.78-1.00). The conserved light response of LOV2 therefore operates within the fusion.

**Figure 5.**
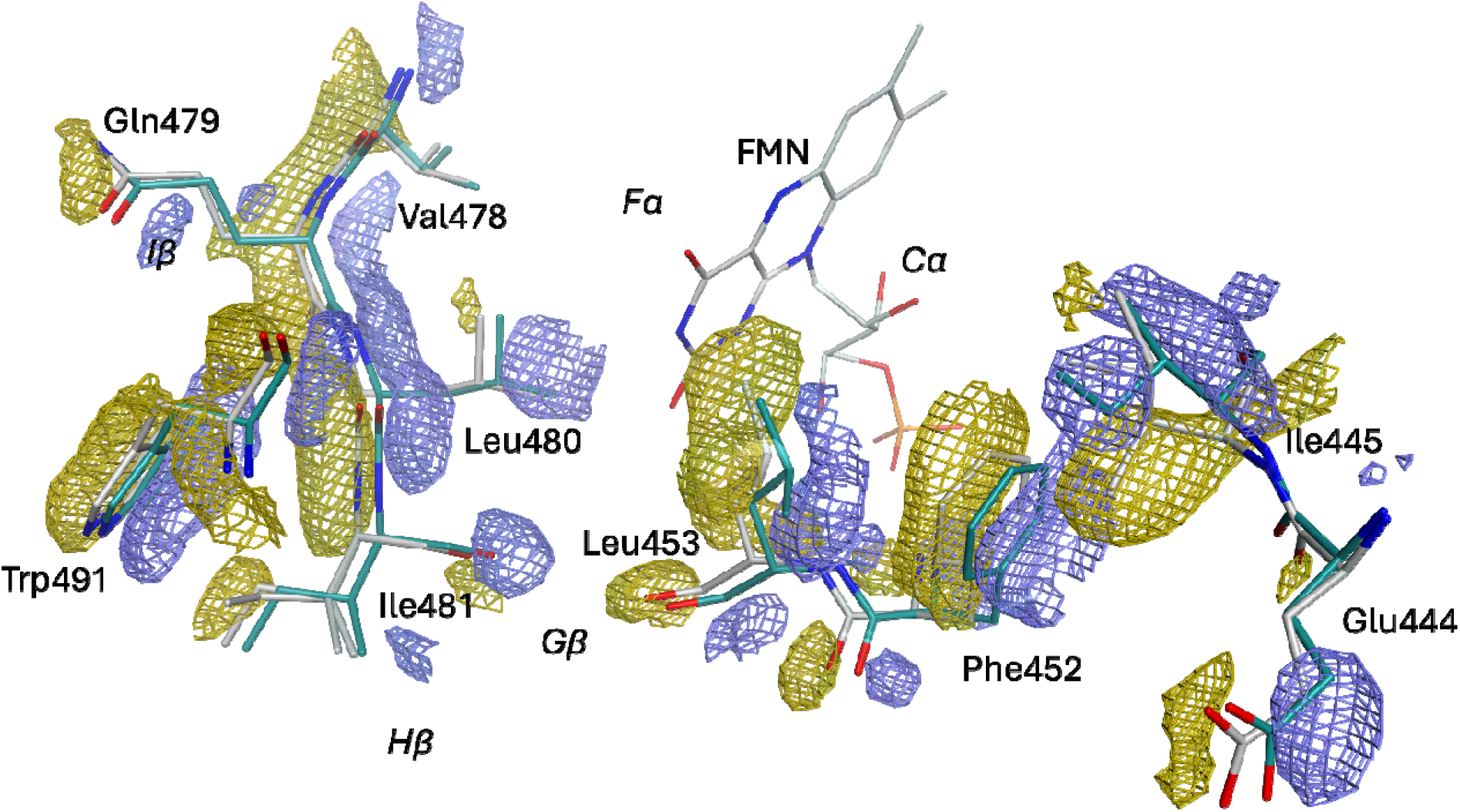
Light-induced structural changes beyond the FMN binding site of AsLOV2-Rac1. The F_light_ − F_dark_ difference electron density map is contoured at ±3.0 σ, with positive and negative features represented as blue and yellow meshes, respectively. Residues undergoing light-induced rearrangements are shown as sticks colored grey (dark state) and deep teal (light state).

### Gln513 and Asn414

The conserved glutamine of LOV domains (Gln513 in AsLOV2) has been studied extensively for its role in signal transduction (Nash *et al*., 2008; Crosson & Moffat, 2001, 2002; Dittrich *et al*., 2005; Nozaki *et al*., 2004; Fedorov *et al*., 2003; Arshi *et al*., 2023; Ganguly *et al*., 2017; Freddolino *et al*., 2006). In the dark it forms a hydrogen bond with FMN O4, and a flip or rotation of its amide has been proposed to let it hydrogen-bond to the newly protonated N5 in the adduct state. This flip was modelled for 80 % of the molecules in the 1.09 Å freeze-trapped light-state structure of isolated AsLOV2 (PDB 7PGY) on the basis of chemical reasoning and spectroscopic evidence (Dietler *et al*., 2022), whereas the earlier cryo-trapped light state 2V0W retains the dark orientation (Halavaty & Moffat, 2007). In our light state, Gln513 shows paired features (+6.0/-5.8 σ) that indicate a small local rearrangement, but the amide keeps its dark-state position, with NE2 3.3 Å from FMN O4 (3.15 Å in the dark) and 3.6 Å from N5 (Figure 6(a)). Because a 180° flip exchanges OE1 and NE2 at almost the same positions, we compared both assignments with density tests calibrated on amides of known orientation (Figures S1 and S2). At 1.8 Å, only the B-factor test discriminates (10 of 11 calibration amides), and it favours the dark orientation. The calibrated contrast test also favours the dark orientation in the 1.09 Å data of 7PGY and the 1.55 Å room-temperature data of 2V1B. A flip in our structure would form no hydrogen bond with the protonated N5, and the equivalent glutamine (Gln489) was also modelled in the dark orientation in room-temperature light states of the LOV2 domain of A. thaliana phototropin 2 (Aumonier *et al*., 2020, 2022). We therefore kept the dark orientation, while noting that it cannot be determined unambiguously at this resolution. Crystallographic and molecular dynamics (MD) studies indicate that this glutamine adopts several side-chain conformations whose populations differ between dark and light states (Freddolino *et al*., 2006; Iuliano *et al*., 2020; Kalvaitis *et al*., 2019), and several LOV receptors retain signalling activity without it, although often with reduced efficiency (Dietler *et al*., 2022). MD simulations of PA-Rac1 proposed that the reoriented Gln513 forms a hydrogen bond with Asn492 (Peter *et al*., 2012a,b); this bond is absent from both our structures (5.0 Å).

**Figure 6.**
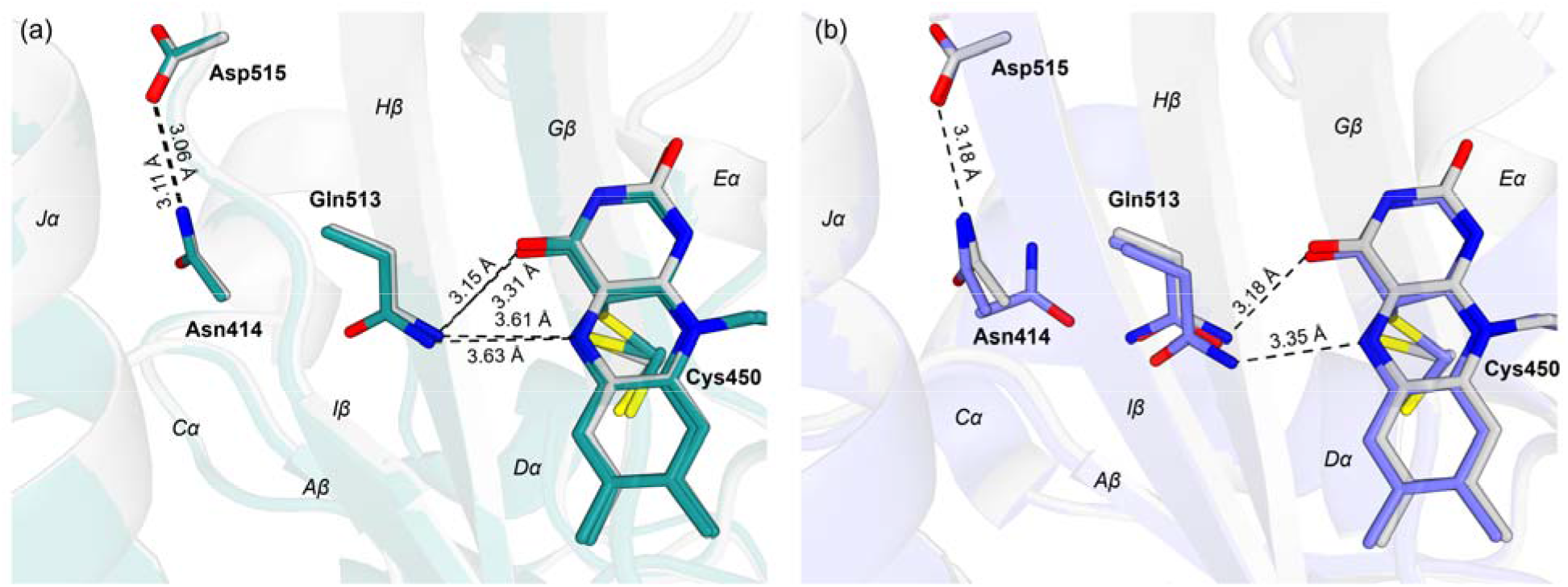
The FMN binding pocket in (a) the dark (grey) and light (deep teal) structures of AsLOV2-Rac1 and (b) the room-temperature dark (grey; PDB: 2V1A) and light (slate blue; PDB: 2V1B) structures of the isolated AsLOV2 domain. Dashed lines show key distances (Å). In (a), 3.06, 3.15 and 3.61 Å refer to the dark state and 3.11, 3.31 and 3.63 Å to the light state; all modelled conformers of Cys450 and FMN are shown, and light-state distances are measured to the adduct conformer (occupancy 0.78). In (b), the 3.18 Å distances are from 2V1A and 3.35 Å from 2V1B (Gln513 conformer A).

In the wild-type light states of the isolated domain, Asn414 is rotated in 60-100 % of the molecules, depending on the structure, and loses its hydrogen bond to Asp515 (Halavaty & Moffat, 2007; Dietler *et al*., 2022) (Figure 6(b)), and in 7PGY the rotated Asn414 stabilizes the flipped Gln513. In PA-Rac1, Asn414 keeps a single orientation and its 3.1 Å hydrogen bond to Asp515 in both states: polder omit maps show density only for the dark-state rotamer, a two-conformer refinement converges to 0.09 for the rotated conformer, and the difference map shows no positive peak at the rotated position (Figure S3), whereas the published rotations are well supported by their own data (5-15 σ). The segment between AZα and Aβ (residues 411-414), which shifts by 0.6-1.2 Å with the rotation in the isolated domain, moves by at most 0.14 Å. In the fusion, the AZα segment packs against Rac1 of the same polypeptide (Leu408 N and Glu409 Oε1 to Asn595, Leu404 to Lys592), and Rac1 fills the position that AZα occupies in the isolated domain (Figure 1(b)). This anchoring may suppress the Asn414 rotation, although intramolecular and lattice contributions cannot be separated in this crystal form.

### The Jα helix and Rac1

NMR and mutagenesis of isolated AsLOV2 showed that Jα packs against the PAS core and that mutations on its hydrophobic face (Ile532Glu, Ala536Glu and Ile539Glu) disrupt this interaction and unfold Jα (Harper *et al*., 2003, 2004). In our light state the difference density around these residues is weak (at most about 4 σ), and the most pronounced Jα features (+4.6/-4.7 σ) lie near Asp522 at its N-terminal end (Figure 7). They arise mainly from a small rigid-body rotation of LOV2 (below): relative to the LOV2 core, the Iβ-Jα junction moves by only 0.07 Å and the Jα axis turns by 0.3°, compared with 0.65-1.02° in the isolated-domain light states. Features of similar magnitude at the C-terminal end of the AZα helix (Ile411, +4.5 σ) and in the adjoining Aβ strand (Phe415-Val416, +5.2/-4.9 σ; Thr418, −4.6 σ) indicate modest backbone adjustments rather than unfolding of the N-terminal cap.

**Figure 7.**
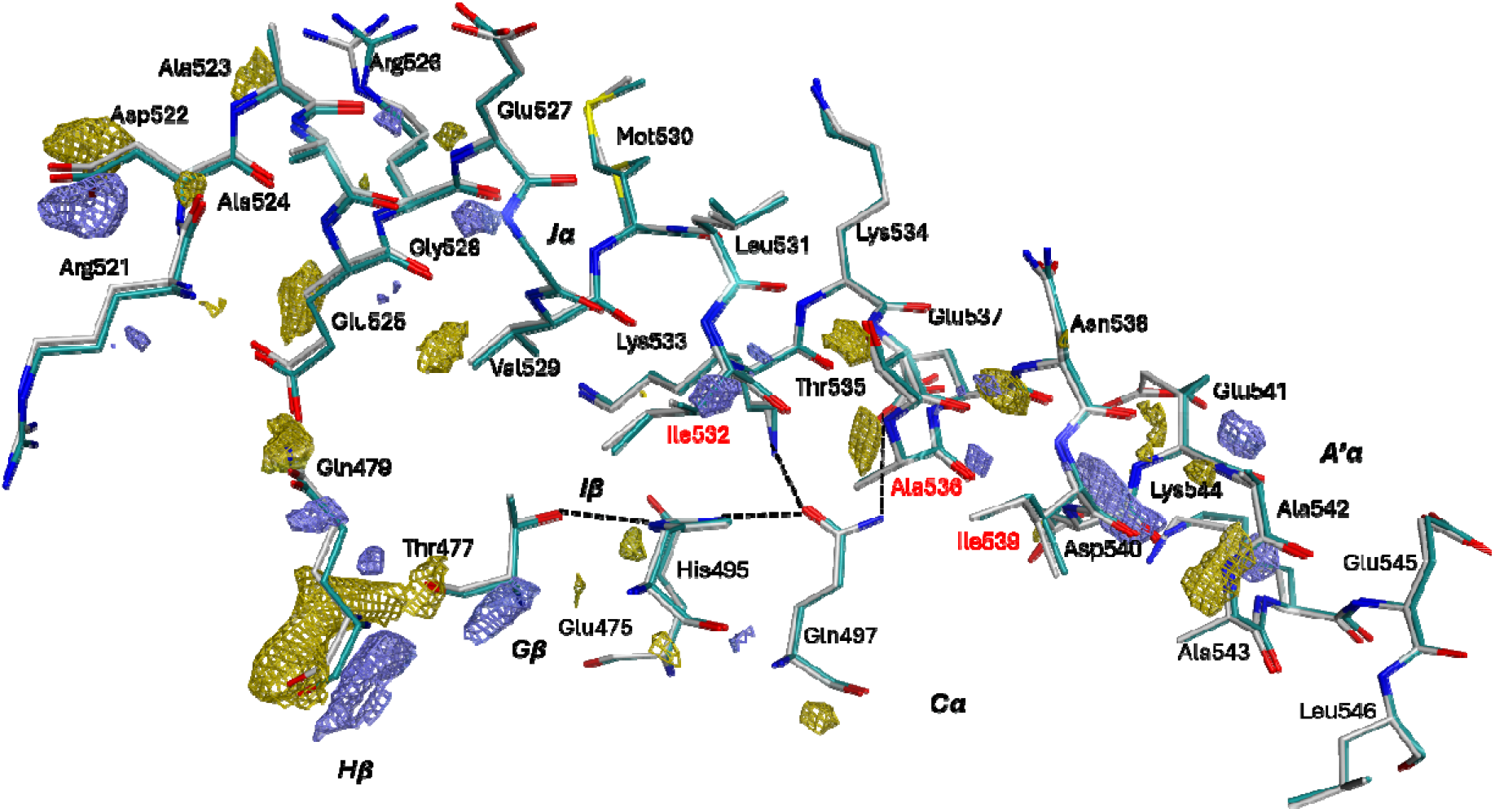
Difference density around the Jα helix of AsLOV2-Rac1. The F_light_ − F_dark_ difference electron density map is contoured at ±3.0 σ, with positive and negative features shown as blue and yellow meshes, respectively. Residues exhibiting positional shifts are shown as grey (dark state) and deep-teal (light state) sticks. Hydrogen bonds are indicated by black dashed lines. Key residues previously identified by NMR as mediating Jα-core interactions are labeled in red.

MD simulations of PA-Rac1 predicted a pathway that starts from hydrogen bonds between the Asp522 amide and Glu525, between the Asp522 and Arg526 side chains and, across the domain interface, between Gln507 and Asp619 and between Glu409 and Asn595; adduct formation was predicted to break these bonds, unwind the N-terminal end of Jα and release Rac1 (Peter *et al*., 2012b). The first three bonds are absent from both our structures and from the cryogenic structure (Wu *et al*., 2009): the Asp522 amide lies 4.1-4.2 Å from Glu525 (3.8 Å in the cryogenic structure), the Asp522 and Arg526 side chains are more than 7 Å apart (6.8 Å), and Gln507 is 6.8 Å from Asp619 (6.7 Å). The crystalline dark state therefore lacks the contacts from which the predicted pathway starts, and the one that is present, Glu409-Asn595, stays intact on illumination (2.8 Å in both states; Figure S4). Likewise, the hydrogen bonds Asp540-Gln497 and Glu518-Gln479 proposed to break in isolated AsLOV2 (Peter *et al*., 2010) are already absent in our dark state, where the alternative contacts Glu537-Gln497 and Glu525-Gln479 are formed and remain intact in the light state (Figure 7).

No local light-induced change is detected in Rac1. Of the 86 unique peaks above 5 σ in the difference map, only two lie in Rac1 (the Cys549 and Cys700 side chains, −5.3 and −5.0 σ), and the nucleotide and Mg^2+^ site show no feature above 4.2 σ. The weaker Rac1 features (at most 4.6 σ) are more numerous than expected from noise and reflect small rigid-body differences between the two structures. A two-body fit to the difference data shows that LOV2 is rotated by 0.73° (bootstrap standard deviation 0.03°) relative to Rac1, about an axis that passes 1.1 Å from the centroid of the LOV2-Rac1 interface and 32 Å from the lattice contacts formed by Glu518 and Asp522, which are preserved (Supporting information). Because the dark and light data come from different crystals (scaled R(F) = 0.15), the data do not show whether this tilt is caused by illumination or by differences between crystals. The largest feature near a Rac1 switch residue lies at the interdomain Ca^2+^ site (+5.9 σ, 0.8 Å from the ion, and −4.6 σ on the Leu446 carbonyl), while the coordination geometry is unchanged (ligand distances change by 0.08 Å or less). Because an isomorphous difference map between two dark-state structures (2WKP and 2WKQ) already shows a feature at this site (Winkler *et al*., 2015), we do not attribute it to illumination.

The Iβ-Jα loop and the N-terminal end of Jα take part in a lattice contact with a symmetry-related Rac1 molecule, through the hydrogen bonds Glu518-Lys696, Asp522 to the main-chain amide of Asp590, Asp522-Arg717 and Asp522-Asp713 (Rac1 Lys153, Asp47, Arg174 and Asp170). Difference features at these residues (for example −4.6 σ at the His519 carbonyl oxygen, measured at the atom) arise mainly from the LOV2 rotation; relative to the LOV2 core, these residues move by 0.15 Å or less. Overall, Rac1 forms more lattice contacts than AsLOV2, which may limit its conformational freedom in the crystal, whereas the isolated AsLOV2 domain forms many more contacts in its own crystals, notably through the N-terminal cap and Jα (Figure S6).

### Matched-temperature comparison of dark and light structures

Cα displacements between our dark and light models, with each domain superposed separately, exceed the 0.30 Å threshold only in the Dα helix (Glu444 and Ile445), at Leu453 next to the FMN, at Ser486 in the Gβ-Hβ loop, at Gln502 and Lys503 in the Hβ-Iβ loop and at the poorly ordered N-terminal residue Leu404 (Figure 8(a)). In Rac1, only Gly573 (Rac1 Gly30, at the start of switch I) in the Pro572-Gly573 loop exceeds the threshold, marginally (0.31 Å); it is modelled in two conformations in both structures and shows no difference-map feature above 5 σ. The Iβ-Jα loop and the Jα helix do not move beyond the threshold. When the same light-state model is compared with the cryogenic dark structure (PDB 2WKP), more residues exceed the threshold: in LOV2, the N-terminal cap (Leu404-Thr407), the Hβ-Iβ loop (Asp501-Gly504) and the C-terminal end of Jα (Ala536, Glu537 and Lys544-Leu546), and, in Rac1, the Pro572-Gly573 loop, where the density is weak, and helix α3 (Asp665-Lys675) (Figure 8(b)). Cryogenic structures can stabilize loops and secondary-structure elements that are more flexible at room temperature, and room-temperature serial crystallography studies have documented temperature-dependent differences: active-site regions can be more mobile at room temperature, a flexible loop can instead be better ordered, ligand binding can differ between temperatures, and active-site conformations can be missing from cryogenic structures (Dunge *et al*., 2024; Günther *et al*., 2025). Comparing structures recorded at the same temperature minimizes these differences and better isolates the light-induced motions.

**Figure 8.**
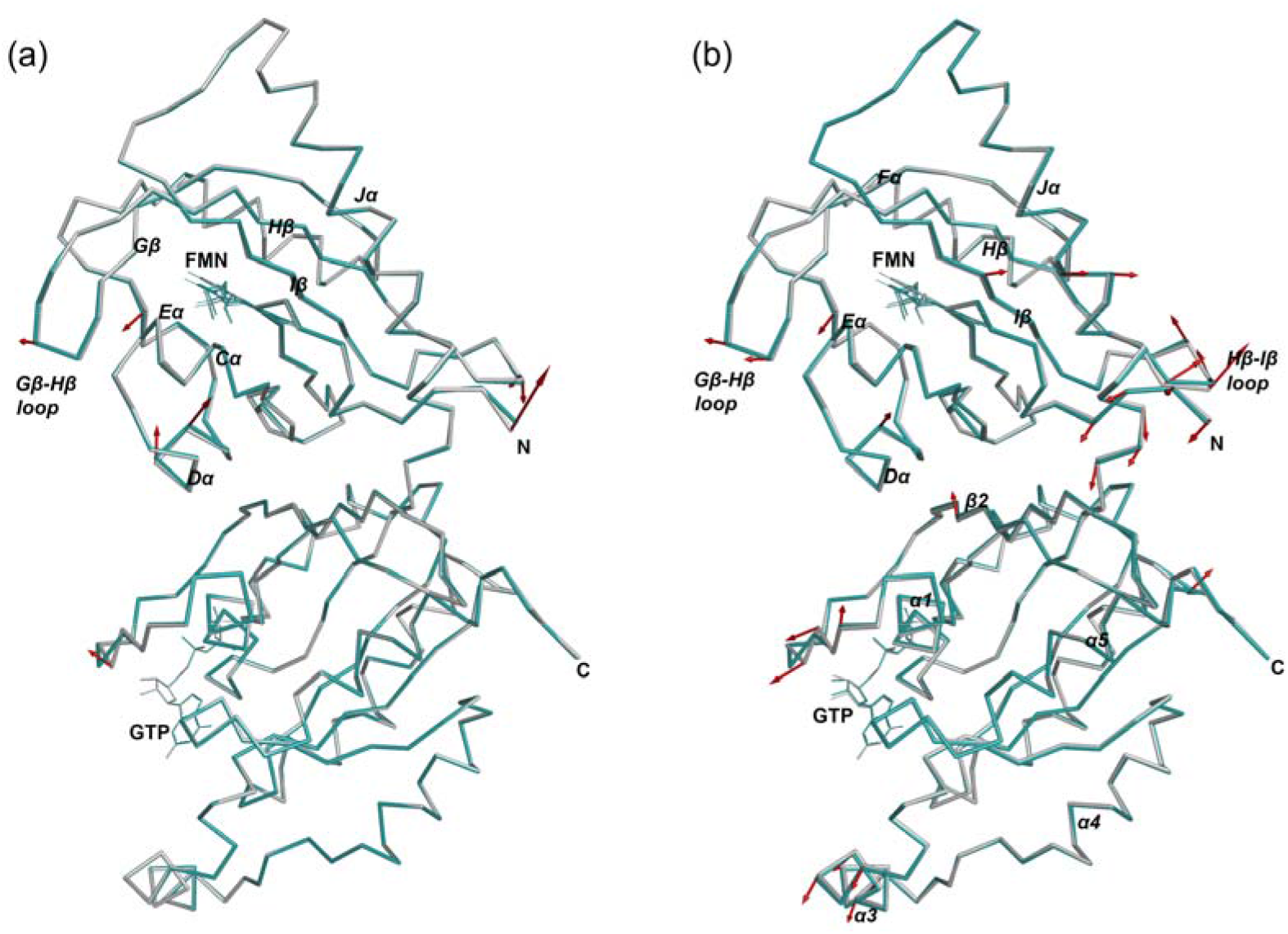
Cα displacements between superposed models of AsLOV2-Rac1. (a) Dark room-temperature (RT) structure (grey) and light-state RT structure (deep teal). (b) Dark cryogenic structure (grey; PDB: 2WKP) and light-state RT structure (deep teal). In each panel, the reference model was superposed onto the light-state model separately for each domain, on the Cα atoms of the LOV2 core (residues 404-521; r.m.s.d. 0.18 Å in (a) and 0.24 Å in (b)) and of Rac1 (residues 547-718; 0.10 and 0.19 Å), and each domain of the reference is drawn from its own superposition. Red arrows start at the Cα atom of the reference model and point towards the light-state position for every Cα that moves by 0.30 Å or more; arrow lengths are eight times the displacement. For residues with alternate conformations, the closest pair of conformers was compared. Arrows were generated with a PyMOL script. Models are shown as Cα traces, with FMN and GTP as thin sticks, in the same orientation in both panels.

Comparisons with the cryogenic and room-temperature structures of the isolated AsLOV2 domain show no global rearrangement; the largest displacements, in the Hβ-Iβ and Gβ-Hβ loops, the Dα and Jα helices and the termini, mainly reflect intrinsic flexibility, domain isolation and crystal packing (Figure S7). The Hβ-Iβ loop is also the most flexible region of the phototropin LOV2 domain in MD simulations (Freddolino *et al*., 2006). The equivalent Gβ-Hβ hairpin of the LOV domain of Chlamydomonas reinhardtii phototropin 1 (Lys92-Thr95) shifts in a similar direction, but by more (Cα shifts of 0.42-0.45 Å after superposition on the LOV core, compared with 0.31 Å at Ser486 here), between the dark and 92.5 ms structures obtained by time-resolved serial synchrotron crystallography (Gotthard, Mous *et al*., 2024), where this hairpin forms a crystal contact (Figure S8).

### Why the Jα helix stays folded in the crystal

In solution, illumination of isolated AsLOV2 unfolds much of Jα. Far-UV circular dichroism reports a loss of about 30 % of the 222 nm signal (Zayner *et al*., 2012), time-resolved infrared spectroscopy resolves Jα unfolding into a biphasic process (Konold *et al*., 2016), and time-resolved X-ray solution scattering shows the AZα and Jα helices unfolding within microseconds (Kim *et al*., 2024). In our light-state structure, the adduct forms and the conserved LOV2 response reaches Dα, Gβ and the Gβ-Hβ loop, but Jα and the N-terminal cap stay folded and no local change is detected in Rac1. These data cannot exclude transient or low-population signalling states, and several factors may contribute to the modest response. First, the energetic driving force may be small. The PAK1 CRIB domain binds the dark-state mutant of PA-Rac1 (C450A) with a K_d o_f 2.3-3.1 µM and the lit-state mutant (I539E) with a K_d o_f 220-280 nM (Wu *et al*., 2009). This 10-fold difference corresponds to only about 1.4 kcal/mol (about 2.3 kT at room temperature). Because the two mutants only mimic the dark and lit states, this is an indirect estimate, but it suggests that the light state may still populate an ensemble close to the dark-state geometry. Second, solution data indicate that activation of PA-Rac1 shifts a pre-existing equilibrium rather than acting as a deterministic switch: hydrogen-deuterium exchange shows caged and unstructured Jα states co-existing even in the dark, and illumination shifts this balance without fully separating LOV2 from Rac1 (Winkler *et al*., 2015). The absence of the Asn414 rotation may also contribute, although this link is speculative: in isolated AsLOV2, MD simulations combined with time-resolved infrared and NMR spectroscopy predict that a transient Gln513-Asn414 hydrogen bond couples the adduct to Jα destabilization, and the N414A mutation partially unfolds Jα already in the dark (Iuliano *et al*., 2020). Third, the crystal may stabilize the caged state, through the lattice contact at the Iβ-Jα loop and the N-terminal end of Jα and through the intramolecular Ca^2+^ bridge between the Dα-Eα segment and switch I, which stabilizes the domain interface in solution (Winkler *et al*., 2015); Ca^2+^ is present at 50 mM or more in the crystallization medium (150 mM calcium acetate in the precipitant solution), far above its dissociation constant of about 0.46 mM. These are plausible, not mutually exclusive explanations rather than established mechanisms, and the present data cannot separate them.

### Conclusions

Room-temperature SFX on SOS chips, combined with repeated pump-laser excitation, gave dark and light-state structures of PA-Rac1 at 1.8 Å with about 80 % adduct, from fragments of large rod-shaped crystals, a morphology that complicates conventional serial crystallography. The conserved LOV2 light response, including the Ile445 rotamer switch, operates within the fusion, whereas the Asn414 rotation of the isolated domain is absent. Apart from the poorly ordered N-terminal residue and a marginal shift in one Rac1 loop, Cα displacements above 0.3 Å between dark and light models recorded at the same temperature are confined to the chromophore environment, the Dα helix and the Gβ-Hβ and Hβ-Iβ loops, whereas comparison with a cryogenic reference can exaggerate the apparent changes. In the crystal, the Jα helix does not unfold, and the GTP-bound Rac1 domain, its Mg^2+^ site and its switch regions show no detectable light-induced change, although transient or low-population states cannot be excluded. This is consistent with the small difference in effector affinity between dark- and lit-state mutants and with the pre-existing conformational equilibrium seen in solution; lattice contacts and the interdomain Ca^2+^ ion may contribute further. To our knowledge, this is the first light-state structure of PA-Rac1 or of any AsLOV2 fusion and the first AsLOV2 light state in which the thioether adduct is not compromised by X-ray radiolysis. SOS chips have been used for SFX and time-resolved pump-probe measurements (Doak *et al*., 2018, 2024; Gorel *et al*., 2025; Barends *et al*., 2026); the present work shows that they also allow a light state to be accumulated by repeated pump excitation during rastering. Because this light state was accumulated rather than resolved in time, it shows the response once the adduct has formed; following the propagation of the signal from the chromophore to Jα and Rac1 may benefit from time-resolved experiments. More generally, light-induced structural changes in photoreceptors are best assessed against a dark reference recorded at the same temperature.

## Author contributions

P.N. conceived and coordinated the project. M.P. and A.S. prepared the samples. J.H.B., M.K. and M.V.A. set up and operated the beamline. T.D.K. loaded the SOS chips. M.K.S., S.M., A.K., A.A., R.K., B.O., M.K. and P.N. collected the X-ray data. M.K.S. and G.G. processed the data. M.W. and P.N. analysed the data. M.W., P.N. and M.P. wrote the original draft. All authors reviewed and approved the manuscript.

## Acknowledgements

We acknowledge the Paul Scherrer Institute, Villigen, Switzerland, for provision of free-electron laser beamtime at the Cristallina beamline of SwissFEL (proposal No. 20240471). Claude (Anthropic), an AI assistant, was used for language editing, reference management, structural analyses and checks, and the preparation of Figures 6, 8, S1-S3, S7 and S8 from the authors’ data and PyMOL scripts; the authors checked all AI-assisted text, analyses and figures.

## Data availability

Atomic coordinates and structure factors of the dark and light-state structures have been deposited in the Protein Data Bank under accession codes 35HK (dark) and 35HL (light).

## Supporting information

### Construct sequence

The crystallized construct (334 residues) consists of an N-terminal tag (MGGSHHHHHHHHGS, not removed), residues 404-546 of A. sativa phototropin 1 and residues 4-180 of human Rac1 carrying the Q61L, E91H and N92H substitutions. Apart from the tag (MRGS followed by six histidines in the original construct), the sequence is identical to that of PDB entry 2WKP. Model residues are numbered from Leu404 of phototropin 1, so that Rac1 residue n is residue n + 543 of the model; the tag is disordered in both structures.

MGGSHHHHHH HHGSLATTLE RIEKNFVITD PRLPDNPIIF ASDSFLQLTE YSREEILGRN CRFLQGPETD

RATVRKIRDA IDNQTEVTVQ LINYTKSGKK FWNLFHLQPM RDQKGDVQYF IGVQLDGTEH VRDAAEREGV

MLIKKTAENI DEAAKELIKC VVVGDGAVGK TCLLISYTTN AFPGEYIPTV FDNYSANVMV DGKPVNLGLW

DTAGLEDYDR LRPLSYPQTD VFLICFSLVS PASFHHVRAK WYPEVRHHCP NTPIILVGTK LDLRDDKDTI

EKLKEKKLTP ITYPQGLAMA KEIGAVKYLE CSALTQRGLK TVFDEAIRAV LCPP

### Comparison with published structures

Coordinates and structure factors of the isolated AsLOV2 dark and light states (PDB entries 2V0U, 2V0W, 2V1A and 2V1B, Halavaty & Moffat (2007); wild-type 7PGX and 7PGY and Q513L 7PGZ and 7PH0, Dietler et al. (2022)) and of PA-Rac1 (2WKP, 2WKQ and 2WKR, Wu et al. (2009)) were taken from the PDB, and R factors and σ_A-_weighted 2mF_o-_DF_c a_nd mF_o-_DF_c m_aps were recalculated for all entries with one protocol in cctbx, as distributed with Phenix (Liebschner *et al*., 2019); for the 2007 entries, which were deposited with residual B factors, the TLS contribution was added back first. All structures were superposed on our dark model using the Cα atoms of residues 415-520, and displacements were considered significant when they exceeded three times the combined diffraction-component precision index of the two structures. The orientation of the Gln513 amide was tested by refining both assignments with otherwise identical protocols (Phenix 1.19.2) and by calibrating B-factor, density-contrast and difference-density tests on asparagine and glutamine amides whose orientation is fixed by unambiguous hydrogen bonds in the same structure (Figures S1 and S2). A rotated Asn414 conformer was tested by two-conformer occupancy refinement and polder omit maps (Liebschner *et al*., 2017) (Figure S3. Displacement arrows (Figures 8, S7 and S8) were generated in PyMOL with a custom script, with arrow lengths proportional to the Cα displacements (scale factors in the figure legends).

### Rigid-body analysis of the difference data

Isomorphous difference amplitudes F(light) − k(s)F(dark), with k(s) refined in 20 resolution shells of equal reflection count, were calculated from amplitudes obtained by refining both datasets with servalcat (Yamashita *et al*., 2021, 2023) from the same starting model, and phased with the dark model. Model difference amplitudes for rigid-body rotations and translations of LOV2 (residues 404-521) and Rac1 (residues 547-718, with GTP and Mg^2+^) were fitted to these data in real space between 2.5 and 8 Å, within masks around each domain that excluded the chromophore pocket, residues with local light-induced changes, the Ca^2+^ site and the nucleotide-binding regions; the tilt is the rotation of LOV2 relative to Rac1. Uncertainties were estimated from 200 bootstrap replicates of the reflections, and null distributions from difference data with randomized signs and from noise matched to the measurement errors. Calculations used gemmi (Wojdyr, 2022) and custom Python scripts. The 0.73° rotation of LOV2 relative to Rac1 lies more than 30 standard deviations above the null fits, and no lattice-contact distance changes by more than 0.09 Å in the fit, so LOV2 pivots on Rac1 while these contacts are preserved. In addition, the whole molecule is rotated by 0.20° in the crystal and Rac1 is displaced by 0.12 Å. Both datasets contain GTP (γ-phosphate occupancy 0.98 and 0.99), so no nucleotide change contributes.

### Cys648 modification

The covalent modification of Cys648 (Rac1 Cys105) gives polder omit peaks of 27.6 and 24.7 σ in the two datasets, 2.2 Å from SG, with three weaker outer lobes, and was modelled as S-sulfocysteine. Arsenic adducts derived from the cacodylate in the crystallization solution are disfavoured by their refined B factors. Additional density at this position was also noted by Wu et al. (2009) in the cryogenic PA-Rac1 structure 2WKP, grown in the presence of cacodylate, and tentatively attributed to a modification by dithioerythritol; in maps calculated from the deposited data, it is absent from 2WKQ and at most weak in 2WKR (about 3 σ), both grown without cacodylate.

### Difference maps of the isolated domain and crystal packing

In an isomorphous light-minus-dark difference map of the room-temperature structures of the isolated AsLOV2 domain (PDB entries 2V1B and 2V1A; Fo-Fo with resolution-shell scaling and phases from 2V1A, 1.65 Å), the features beyond the chromophore are weaker than in our map, although the adduct gives a strong signal (+15.7 σ at Cys450): peaks at Phe452, Trp491 and the Gβ residues Val478-Ile481 reach 3.3-4.8 σ, compared with 5.0-8.8 σ in our map, and only Leu453 reaches a similar level (+5.4/-4.9 σ). Part of this difference may reflect the lower isomorphism of the two isolated-domain crystals (isomorphous R factor 0.30). In the crystals of the isolated AsLOV2 domain, lattice contacts involve the N-terminal cap and Jα, which may hinder the light-induced displacement of these elements, as well as the Eα-Fα, Gβ-Hβ and Hβ-Iβ loops, strand Bβ and helix Dα; the latter contacts are absent in the AsLOV2-Rac1 crystals, in which the main AsLOV2 lattice contact involves the Iβ-Jα loop and the N-terminal end of Jα (Figure S6). In PA-Rac1, the Rac1 regions in lattice contacts include the segment containing both 3_10 h_elices, the α1, α3 and α5 helices, the β2-β3 and β6-α5 loops and the entire β6 strand.

## Supplementary figures

**Figure S1.**
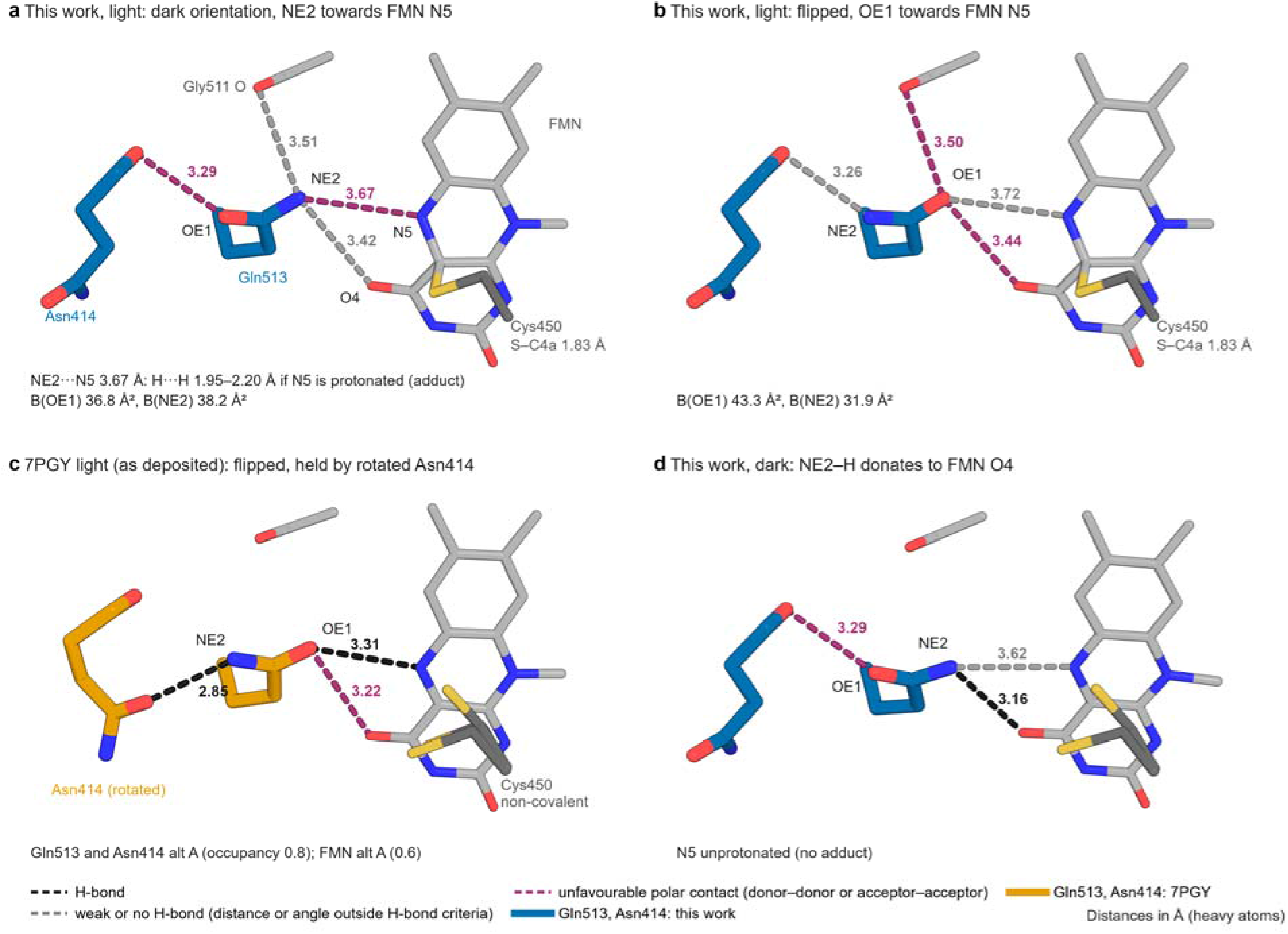
Hydrogen bonding of the two possible Gln513 amide orientations in the adduct state. Gln513, Asn414, the Gly511 carbonyl, the flavin and Cys450 are shown for (a) the light state of this work with the adduct-state Gln513 conformer in the dark orientation, (b) the same model with the amide flipped and refined by an identical protocol, (c) the light state of isolated AsLOV2 at 1.09 Å (PDB 7PGY) as deposited and (d) the dark state of this work. Cys450 is drawn as the thioether adduct bonded to FMN C4a where the model is covalent (a, b); 7PGY was deposited without the bond (c). Dashes join amide heavy atoms to polar neighbours, with distances in Å: black, hydrogen bonds; grey, weak or absent hydrogen bonds; purple, unfavourable contacts between two donors or two acceptors. In (a), with N5 protonated, the NE2-H and N5-H hydrogens come within 1.95-2.20 Å of each other; the flip in (b) removes this contact but forms no hydrogen bond, and it splits the amide B factors (43.3 and 31.9 Å², against 36.8 and 38.2 Å² in (a)), the pattern that marks a wrong O/N assignment in 10 of 11 calibration amides (Figure S2). In 7PGY (c) the flipped amide donates a hydrogen bond to the rotated Asn414, which our light state lacks (Figure S3). R_free_ is 0.1846 for (a) and 0.1847 for (b).

**Figure S2.**
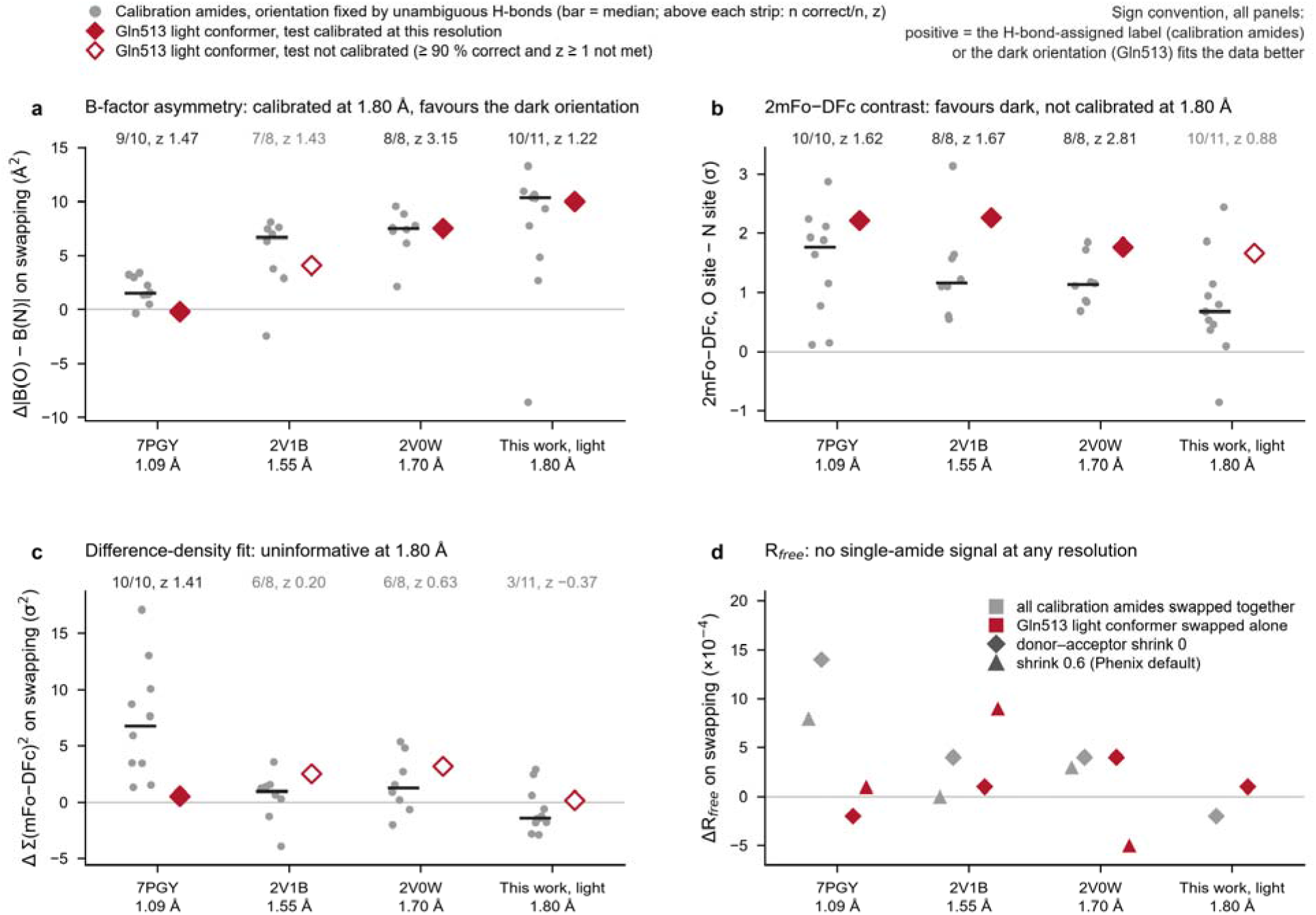
Calibrated density tests for the orientation of the Gln513 amide. Each test was calibrated on asparagine and glutamine side chains whose amide orientation is fixed by unambiguous hydrogen bonds (grey points; bar, median; n = 10 for 7PGY, 8 for 2V1B, 8 for 2V0W and 11 for this work). Values are signed so that a positive value means that the hydrogen-bond assigned label (calibration amides) or the dark orientation (Gln513, crimson diamonds) fits the data better; filled diamonds mark tests that are calibrated at that resolution (at least 90 % of calibration amides correct and z of at least 1). (a) B-factor asymmetry on swapping the O and N labels. (b) 2mF_o_-DF_c_ contrast between the O and N sites. (c) Difference-density fit. (d) Change in R_free_ on swapping. At 1.80 Å only the B-factor test is calibrated (10 of 11 amides), and it places the adduct-state Gln513 in the dark orientation (+10.0 Å², calibration median +10.3 Å²). At 1.09 Å all three density tests are calibrated, and the contrast test favours the dark orientation for the light-state Gln513 conformer of 7PGY (+2.2 σ). At 1.55 Å the contrast test is still calibrated and also favours the dark orientation for the partially flipped Gln513 of 2V1B (+2.3 σ). R_free d_oes not resolve the orientation of a single amide at any of these resolutions. The published entries were re-refined against their deposited structure factors in Phenix 1.19.2; σ is the root-mean-square density of each map over the unit cell.

**Figure S3.**
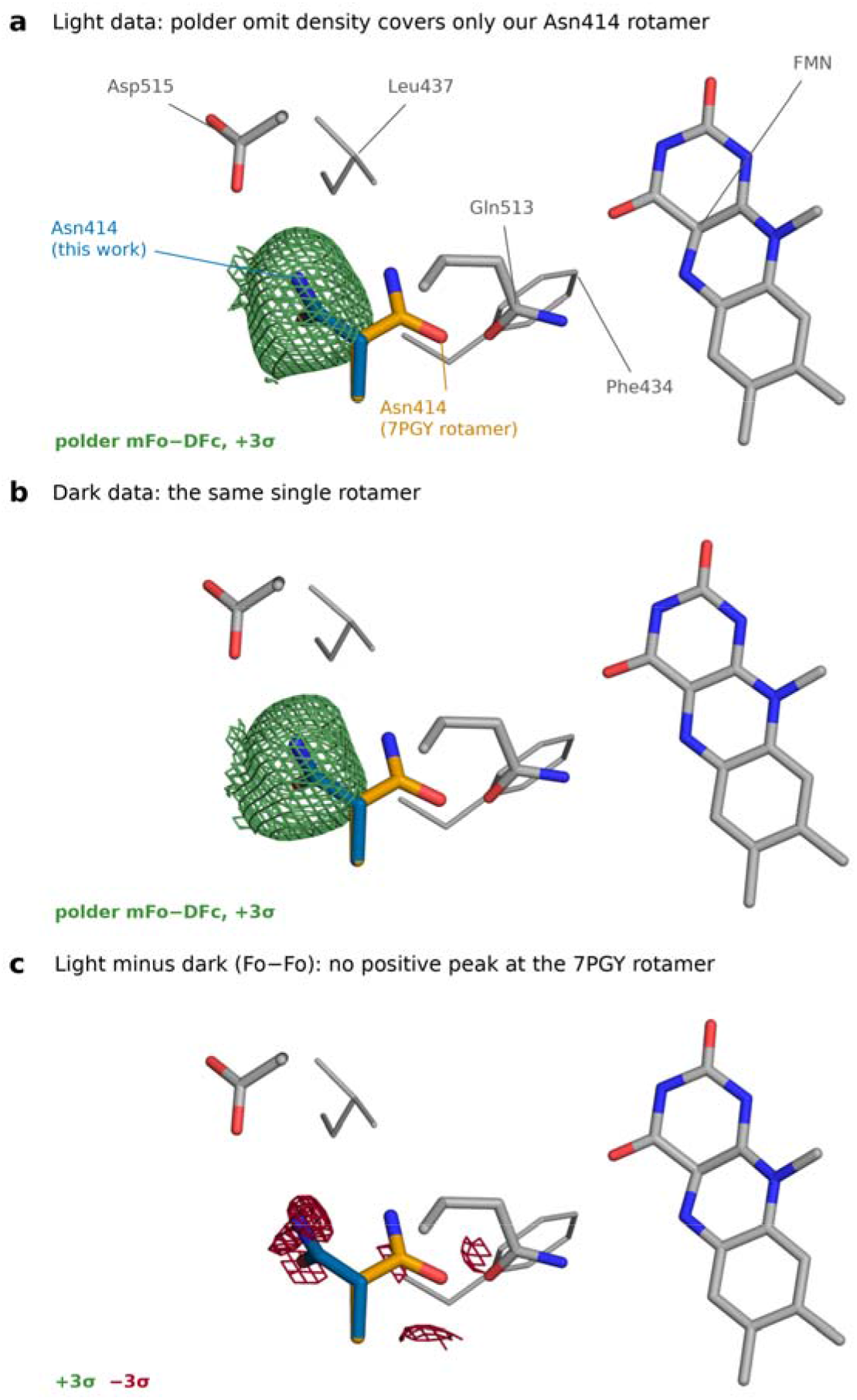
Asn414 keeps its dark-state rotamer in the light state. Asn414, Gln513, Asp515 and the flavin are shown, with Asn414 as modelled in this work (blue; χ1 −57°) and the rotated Asn414 of the isolated-domain light state 7PGY (orange; χ1 −172°) placed on our main chain after superposition on the LOV core (Cα atoms 415-520). (a) Polder omit map of the light data with the Asn414 side chain beyond Cβ omitted, contoured at +3 σ. (b) The same for the dark data. (c) Light-minus-dark difference map contoured at ±3 σ. At the atom positions, the polder map reaches 14.1-16.2 σ at our conformer in the light data (17.9-19.7 σ in the dark data) and −0.5 to 1.8 σ at the rotated conformer; the difference map is −3.5 σ at our ND2 and 1.4-1.9 σ at the rotated conformer, where a rotation in 20 % of the molecules would give about +3 σ. Refinement of the two conformers gave 0.09 for the rotated conformer in the light data and 0.00-0.01 in the dark data. σ is the root-mean-square density of each map over the unit cell.

**Figure S4.**
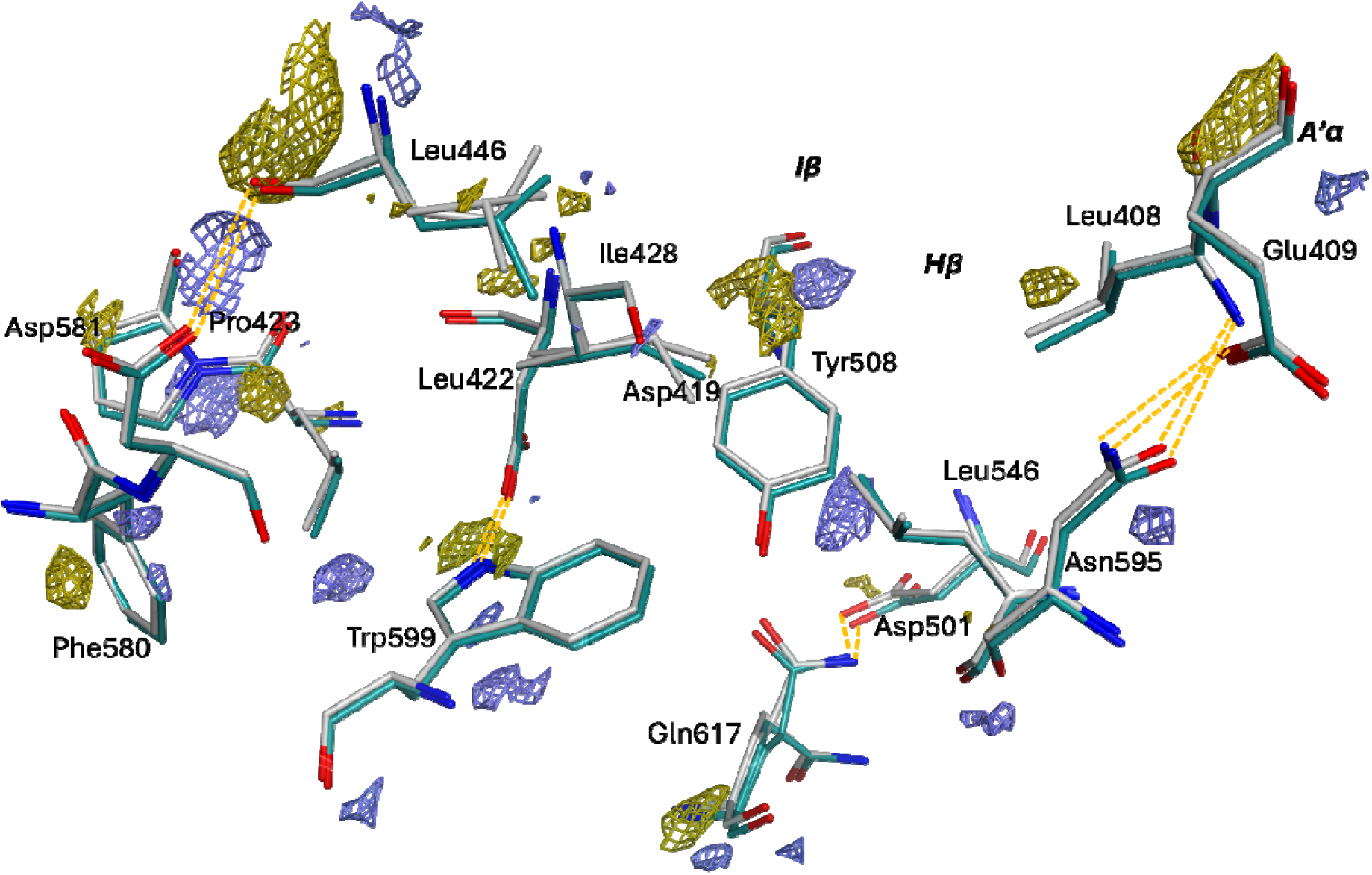
Close-up view of the AsLOV2-Rac1 interface. Residues participating in contacts are shown as grey (dark state) and deep teal (light state) sticks, and hydrogen bonds are indicated with dashed lines. The difference electron density map F_light_ − F_dark_ contoured at ±3.0 σ (positive: blue, negative: yellow) is displayed around these residues, highlighting local structural changes between states.

**Figure S5.**
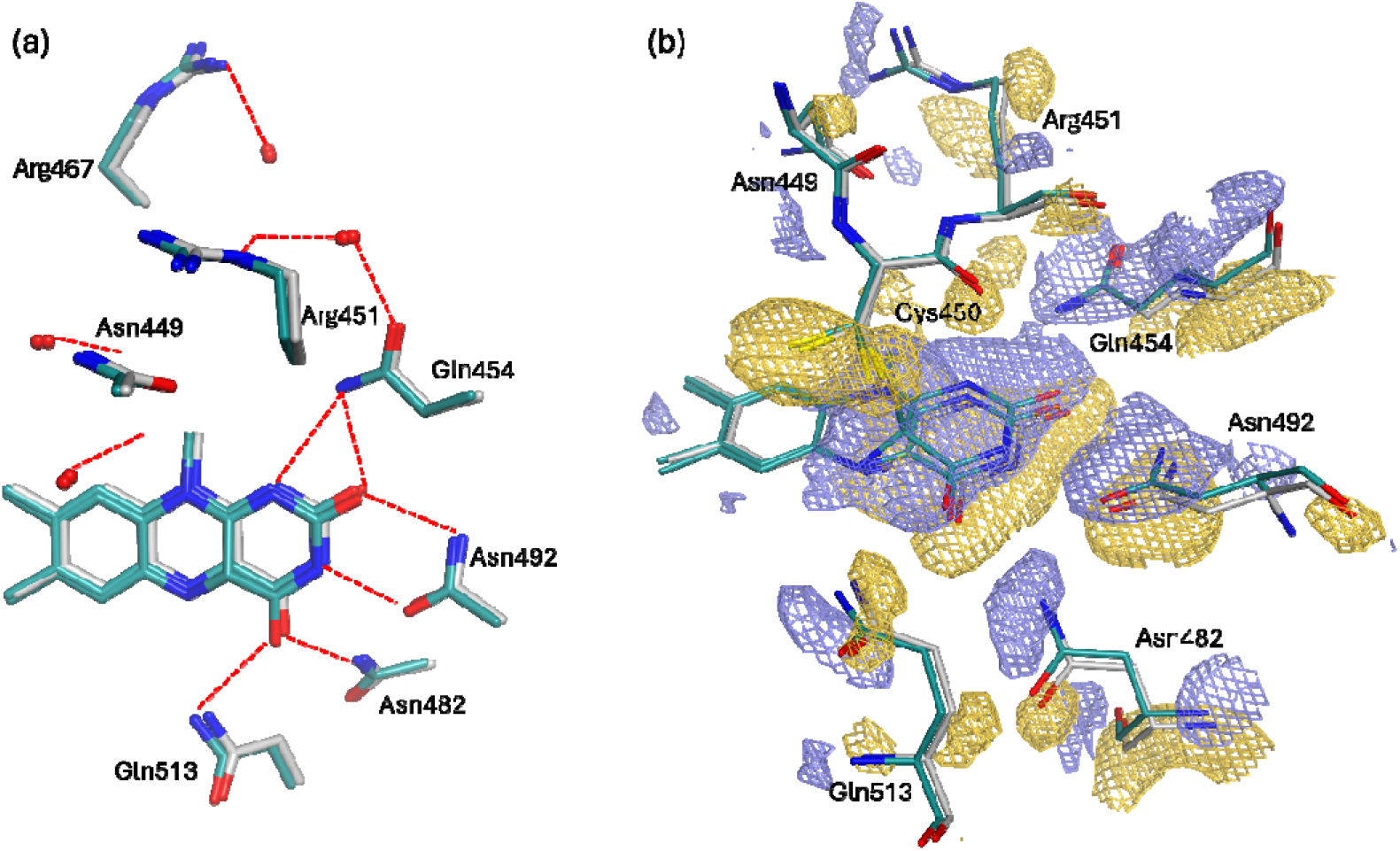
The FMN chromophore binding site upon blue-light activation of AsLOV2-Rac1. (a) Hydrogen-bonding network stabilizing the FMN cofactor in the dark state (grey) and the light state (deep teal). Water molecules are shown as red spheres, and hydrogen bonds as red dashed lines. (b) Light-induced structural rearrangements around the FMN. The F_light_ − F_dark_ difference electron density map is contoured at ±3.0 σ, with positive and negative features displayed as blue and yellow meshes, respectively. Residues exhibiting conformational shifts are shown as sticks colored grey (dark) and deep teal (light).

**Figure S6.**
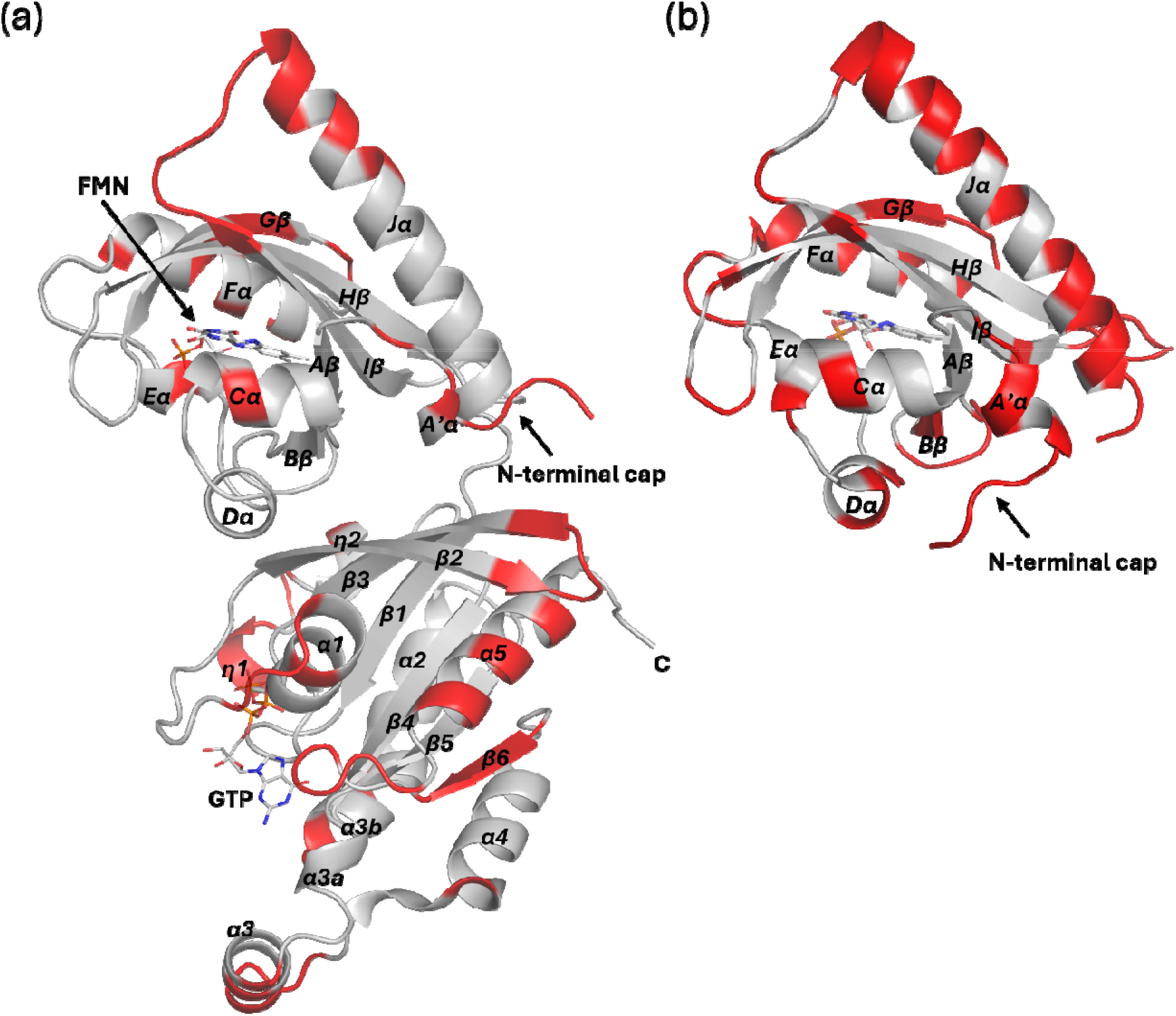
Intermolecular crystal contacts in the structures of the light state of AsLOV2-Rac1. (a) and the room-temperature light state of AsLOV2 (PDB: 2V1B) (b). Residues highlighted in red are within 5 Å of any symmetry mate.

**Figure S7.**
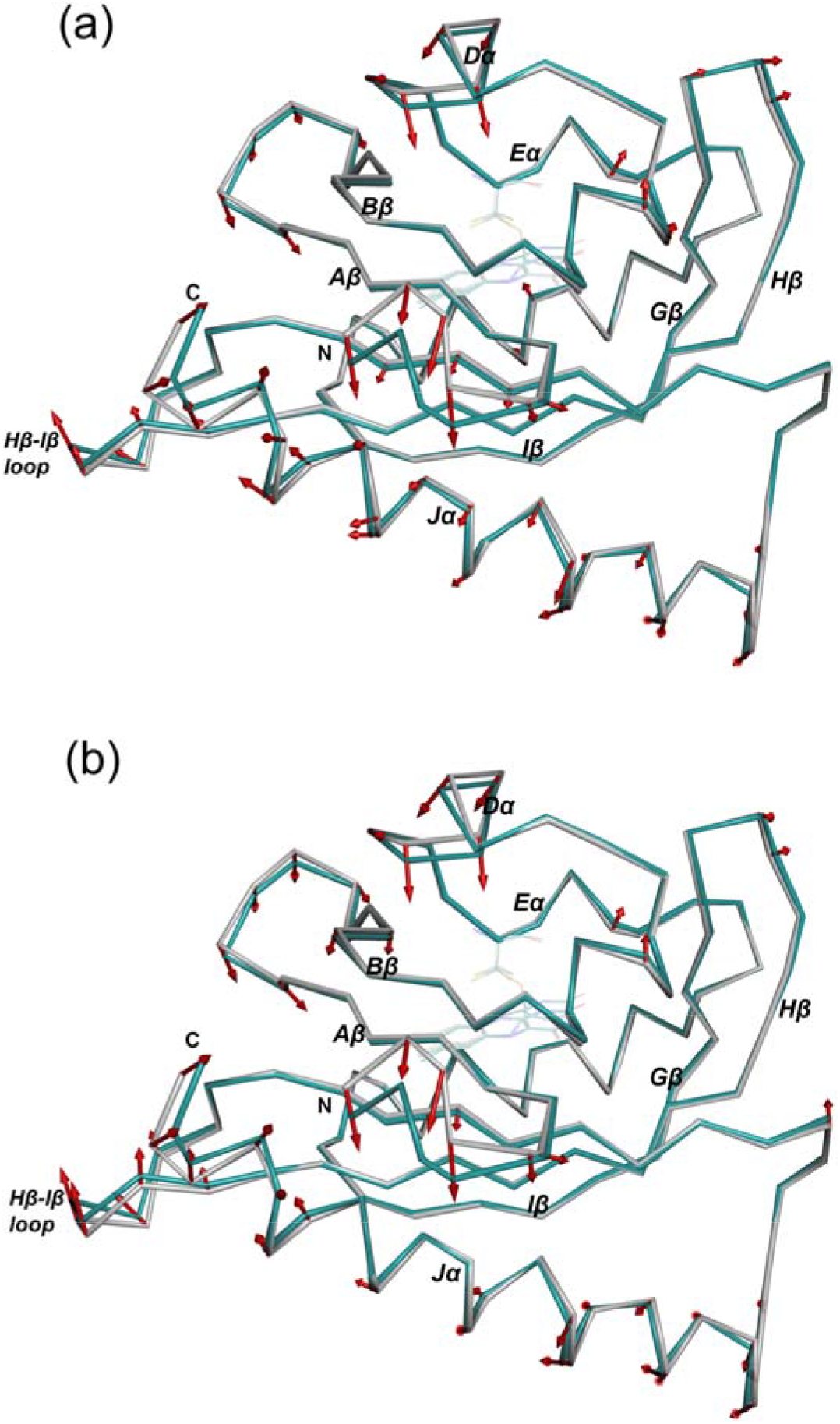
Cα displacements between dark-state structures of the isolated AsLOV2 domain and AsLOV2 in the light-state structure of AsLOV2-Rac1. (a) Dark cryogenic structure of the isolated domain (grey; PDB: 2V0U) and the room-temperature light-state structure (deep teal). (b) Dark room-temperature structure of the isolated domain (grey; PDB: 2V1A) and the light-state structure (deep teal). Residues 408-546 are shown as Cα traces, with Cys450 and the FMN isoalloxazine ring as faint sticks. The structures were superposed on the Cα atoms of residues 415-520 (106 pairs; r.m.s.d. 0.32 Å for 2V0U and 0.36 Å for 2V1A). Red arrows start at the Cα atom of the isolated-domain structure and point towards the light state for every Cα that moves by 0.30 Å or more; arrow lengths are four times the displacement, and displacements larger than 0.8 Å are drawn at 0.8 Å (15 residues in (a) and 22 in (b), including the termini). For residues with alternate conformations, the closest pair of conformers was compared. Arrows were generated with a PyMOL script.

**Figure S8.**
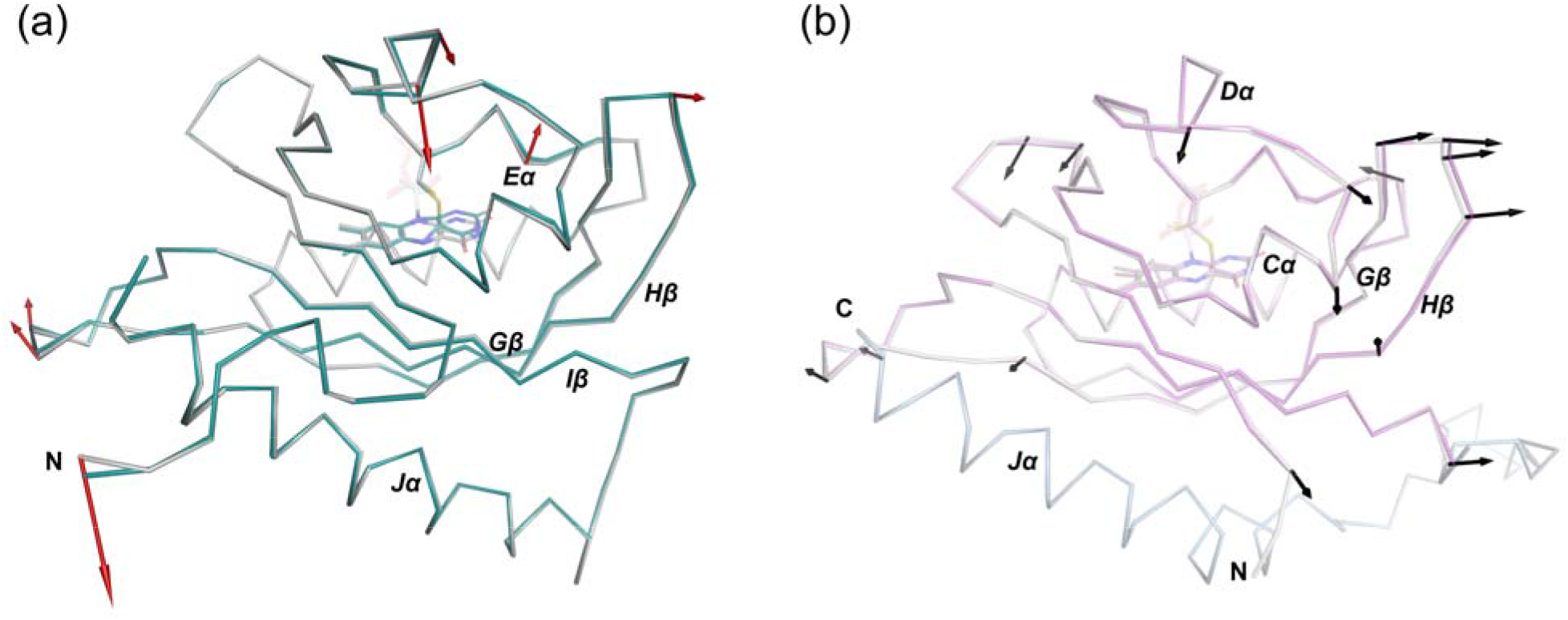
Light-induced Cα displacements in the LOV domains of AsLOV2 and CrPhotLOV1. (a) AsLOV2 in AsLOV2-Rac1 (this work): dark (grey) and light-state (deep teal) room-temperature structures, superposed on the Cα atoms of residues 404-521 (118 pairs; r.m.s.d. 0.18 Å); red arrows. (b) CrPhotLOV1: dark structure (grey; PDB: 8QI9) and light-state structure 92.5 ms (90-95 ms) after illumination (magenta; PDB: 8QIW) from time-resolved serial synchrotron crystallography, superposed on the Cα atoms of residues 19-124 (106 pairs; r.m.s.d. 0.24 Å); black arrows. The Jα helix, which is not modelled in 8QI9 or 8QIW, is shown from the AlphaFold model AF-Q8LPD9-F1 (light blue) after superposition on the same Cα atoms. Arrows start at the dark-state Cα and point towards the light-state position for every Cα that moves by 0.30 Å or more; arrow lengths are eight times the displacement. For residues with alternate conformations, the closest pair of conformers was compared. FMN and the reactive cysteine (Cys450 in (a), Cys57 in (b)) are shown as sticks. The longest arrow in (a) is at the N-terminal residue Leu404.

